# Role of Rv3351c in trafficking *Mycobacterium tuberculosis* bacilli in alveolar epithelial cells and its contribution to disease

**DOI:** 10.1101/2020.12.03.409664

**Authors:** Megan Prescott, Kari Fine-Coulson, Maureen Metcalfe, Tuhina Gupta, Michelle Dookwah, Rebecca Pavlicek, Hind Yahyaoui Azami, Barbara Reaves, Ahmed Hikal, Michael Tiemeyer, Russell Karls, Frederick Quinn

## Abstract

Although interactions with alveolar macrophages have been well characterized for *Mycobacterium tuberculosis*, the roles epithelial cells play during infection and disease development have been less studied. We have previously shown that deletion of gene *rv3351c* reduces *M. tuberculosis* replication in and necrosis of A549 human type II pneumocyte cells. In the present study, we report that *rv3351c* is required for lipid raft aggregation on A549 cell plasma membranes during *M. tuberculosis* infection. Lipid raft aggregation was also induced directly by recombinant Rv3351c protein. A *Δrv3351c* deletion mutant was less effective than wild type *M. tuberculosis* at circumventing phagolysosome fusion in A549 cells as evidenced by increased co-localization with lysosomal markers LAMP-2 and cathepsin-L by the mutant bacilli. These observations indicate a role for Rv3351c in modification of the plasma membrane to facilitate trafficking and survival of *M. tuberculosis* bacilli through alveolar epithelial cells, and support the hypothesis that *M. tuberculosis* has mechanisms to target the alveolar epithelium. Preliminary data also demonstrate that like the type II pneumocyte-targeting *M. tuberculosis* secreted protein heparin-binding filamentous hemagglutinin (HBHA), Rv3351c is detected by the host cellular and humoral immune responses during infection, and may play an important role in mycobacterial dissemination from the lungs.

**Author summary:** *Mycobacterium tuberculosis* is the leading causes of death due to a single infectious agent and many facets regarding the pathogenesis of this organism remain unknown. This facultative intracellular bacterial pathogen often establishes infection through inhalation of the bacilli into the alveoli of the lungs. Interactions with alveolar macrophages have been well characterized and it had been assumed that these interactions with phagocytic cells primarily determine the fate of the disease. However, alveolar epithelial cells, such as type II pneumocytes, play important roles in disease progression of other bacterial and viral respiratory pathogens, which provided the impetus to more-closely examine pneumocyte-*M. tuberculosis* interactions. We describe in this study the role of the *M. tuberculosis rv3351c* gene product in the internalization and survival of this pathogen in human type II pneumocytes. We previously showed that a *Δrv3351c* mutant replicates less efficiently and generates less necrosis than the parental *M. tuberculosis* strain in this cell type. We demonstrate herein that Rv3351c protein induces lipid raft aggregation on the membranes of alveolar epithelial cells and that *M. tuberculosis Δrv3351c* traffics through LAMP-2-labeled endosomes 30% more frequently than the parent strain. This trafficking toward phagolysosomes may underlie the reduced replication and cytotoxicity of the mutant. The role of Rv3351c in trafficking and survival of *M. tuberculosis* bacilli through epithelial cells ultimately resulting in dissemination from the lungs may begin with modifications to the plasma membrane prior to attachment. Such a mechanism of activity suggests Rv3351c as a potential vaccine target to train the host immune system to bind and eliminate the protein before it modulates the alveolar epithelium.

## Introduction

*Mycobacterium tuberculosis*, the causative agent of tuberculosis infects an estimated one-quarter of the world’s population with 60-90% of these individuals potentially harboring latent infection (1,2). To date, no vaccine reproducibly protects against the pulmonary form of the disease in post-adolescents.

The alveolar macrophage is generally believed to control the initial success or failure of *M. tuberculosis* infections. While based in large part on the lack of *in vivo* data from early-stage infection and clinical studies, alveolar epithelial cells (AECs) were not considered important players in *M. tuberculosis* pathogenesis (3,4). However, a growing number of studies suggest they may serve as hosts to this pathogen (5-7). *In vitro* studies have demonstrated that *M. tuberculosis* bacilli can enter and replicate to high numbers in type II pneumocyte cell lines in monolayers or in epithelial/endothelial bilayer systems (6-12); however, *M. tuberculosis* entry into AECs is much slower than observed in macrophages. McDonough and Kress determined that virulent *M. tuberculosis* bacilli can infect both polarized and non-polarized A549 cells (19). An understanding of the mediators and mechanisms responsible for microbial attachment and internalization in these epithelial cells is important. The *M. tuberculosis* heparin-binding hemagglutinin facilitates attachment to sulfated glycoconjugates which are more abundant on AECs than on macrophages, and this protein promotes dissemination of the pathogen from the lungs (9,13). Chitale *et al*. determined that products of the *M. tuberculosis mce* locus promote uptake into human epithelial (HeLa) cells (15). Hsu, *et al*. reported pneumocytes and macrophages infected with a strain of *M. tuberculosis* deficient in the critical virulence factor Early Secretory Antigenic Target-6 (ESAT-6) resulted in the loss of a cytolytic phenotype in both cell types (14). Our laboratory demonstrated that intracellular trafficking patterns for *M. tuberculosis* bacilli in AECs significantly differ from those in macrophages, and that autophagy is a crucial part of this process (16); these observations have been confirmed by other groups (17,18). Taken together, these finding support the hypothesis that cells of the alveolar epithelium can serve as potential hosts for *M. tuberculosis* bacilli, particularly in early stages of infection.

While cell death in macrophages resulting from virulent or avirulent mycobacterial infections can proceed via apoptotic or non-apoptotic pathways, AEC death following *M. tuberculosis* infection is reported to result primarily in necrosis by an undefined mechanism (20-22). The ability of *M. tuberculosis* bacilli to kill type II pneumocytes via this route is potentially an indicator of strain virulence. Our laboratory generated a *M. tuberculosis* mutant lacking the *rv3351c* gene, which like a heparin-binding filamentous hemagglutinin (*hbha*) mutant, is less effective than the virulent parent strain at killing type II pneumocytes (8,13).

The present study examines attachment, internalization and trafficking within type II pneumocytes by *M. tuberculosis* strain Erdman and mutant *Δrv3351c*. In addition, dissemination from the lungs of mice infected with *Δrv3351c* bacilli and humoral immune responses to the Rv3351c protein in sera from human tuberculosis patients is examined. The presented findings shed additional light on the roles of pneumocytes in mycobacterial respiratory disease development and dissemination.

## Results

### Rv3351c induces lipid raft aggregation on epithelial cell plasma membranes

We previously observed that A549 alveolar epithelial cells (AECs) infected with *M. tuberculosis* Erdman bacilli or culture filtrates from those infections added to fresh monolayers induce lipid raft aggregation on the host cell plasma membranes to levels equal or more than lipid raft super-aggregator listeriolysin O (LLO) (9). In the present study, we show that *Δrv3351c* bacilli induce lipid raft aggregation on infected A549 cells at a significantly lower rate than wild-type Erdman bacilli in fluorescent confocal microscopic examinations of *M. tuberculosis-infected* and cholera toxin subunit B-labeled A549 monolayers; cholera toxin binds to lipid raft marker GM1. At six hours post-infection (hpi), 1.5 versus 0.2 lipid rafts per cell, respectively, were observed in cells infected with the wild type versus mutant strain (Figure 1, panels A, B, and F). Addition of recombinant Rv3351c protein alone induced lipid raft aggregtion on A549 cells to a similar extent as addition of listeriolysin O (LLO) protein relative to untreated cells (Figure 1, panel C-F), or cells treated with another recombinant mycobacterial protein (Rv0097) of similar molecular weight (Figure 1, panel G). Additionally, the Rv3351c protein bound to GM1 by thin-layer chromatography overlay analysis (Supplemental Figure 1), suggesting Rv3351c may interact directly with lipid raft components.

**Figure 1.**
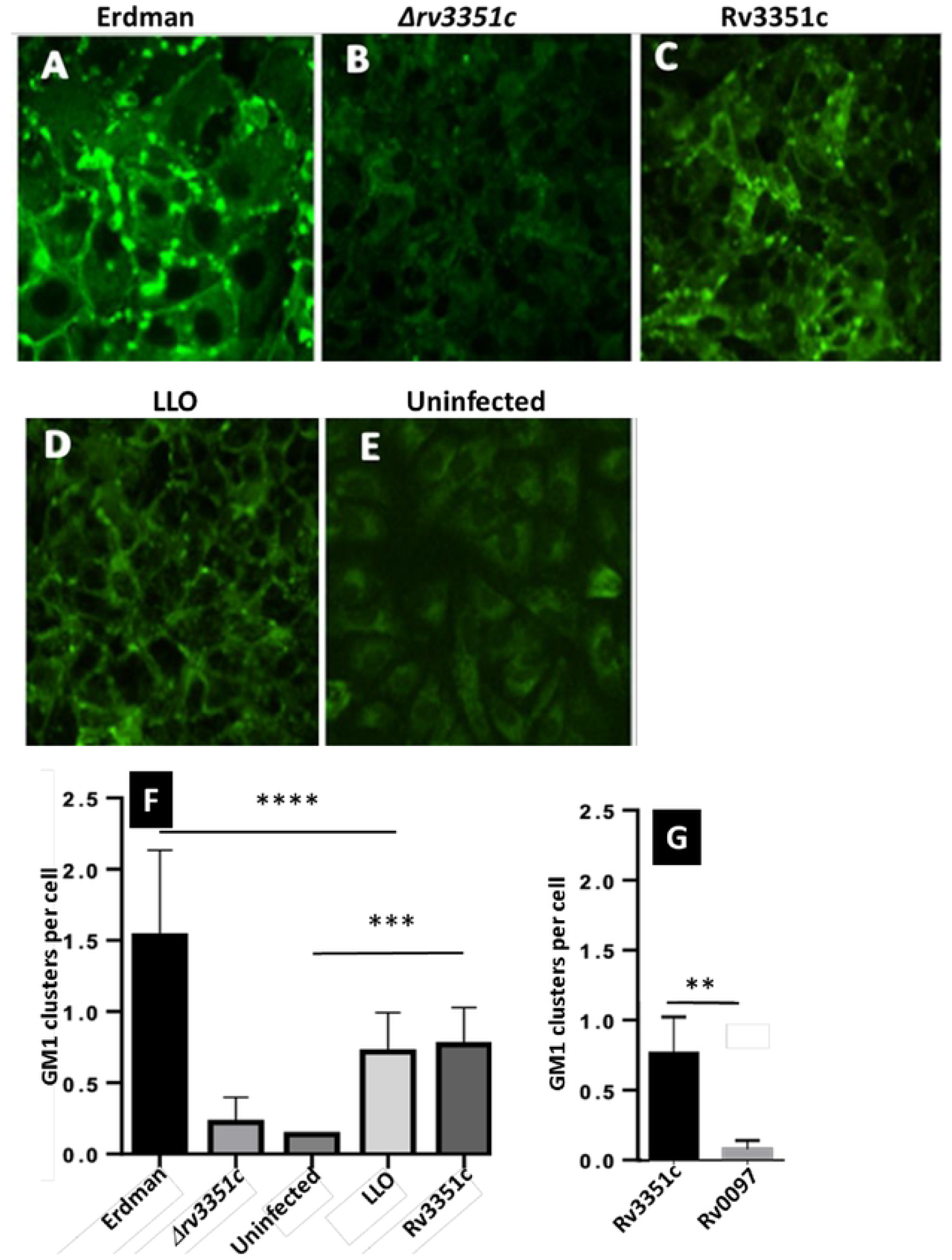
rv3351c induces lipid raft formation in A549 cells. A549 cells infected with MOI = 100 of the indicated *M. tuberculosis* strain were immunostained for lipid raft marker GM1 and examined by confocal microscopy. At 6 hpi, confocal z-stack images showed less GM1 staining (green) of cells infected with (A) *M. tuberculosis* strain Erdman versus (B) *Δrv3351c*. Uninfected control cells treated with (C) Rv3351c protein (2 μg) showed induction of lipid rafts comparable to (D) uninfected cells treated with LLO (2 μg). (E) Untreated control cells showed limited GM1 staining. (F) Quantification of confocal data collected from the experimental groups represented in panels A-E. (G) Counts of GM1 clusters demonstrates differences in lipid raft numbers between A549 cells incubated for 30 minutes with *M. tuberculosis* Rv3351c or control RvOO97 proteins. Lipid rafts defined as GM1 clusters greater than 200 nM in size are represented. Data shown are means of three experiments performed in triplicate, ****p-values: (p<0.001), ***p-values: (p=0.001), **p-values: (p<0.05). Confocal image A was captured at 63x magnification with 3x zoom; all other confocal images were at 63X. Error bars represent one standard deviation.

### Induction of lipid raft aggregation by Rv3351c is essential for *M. tuberculosis* virulence

We previously observed that the aggregation of lipid rafts is critical to *M. tuberculosis* Erdman survival in A549 cells (9). Specifically, disrupting lipid rafts with lipid raft/cholesterol disrupting agent Filipin III decreased the intracellular viability of *M. tuberculosis* bacilli. To determine if the absence of *rv3351c* was responsible for the observed absence of lipid rafts and subsequent loss of virulence in cells infected with the mutant strain, we aggregated lipid rafts with Rv3351c protein before infection with the *Δrv3351c* bacilli. Briefly, A549 monolayers were incubated with 5 μg of recombinant Rv3351c protein, and after 30 minutes, the cells were washed and maintained in cell culture medium for the duration of the experiment. The addition of Rv3351c restored the virulence phenotype of *Δrv3351c* bacilli to wild-type levels over the course of infection as measured by lactate dehydrogenase (LDH) release into the cell culture medium (Figure 2A and B). Lactate dehydrogenase is only released from cells with damaged cell membranes such as those undergoing necrosis or late stage apoptosis. Similar to our previous study, disrupting lipid rafts with Filipin III significantly decreased the rate of host cell killing by Erdman bacteria, but had no significant effect on cells infected with *Δrv3351c* bacilli was observed (Figure 2C and D). Equal amounts (5 μg per well) of Rv3351c or Filipin III were used in this assay and no cytotoxic effects were observed with either at the time points used.

**Figure 2.**
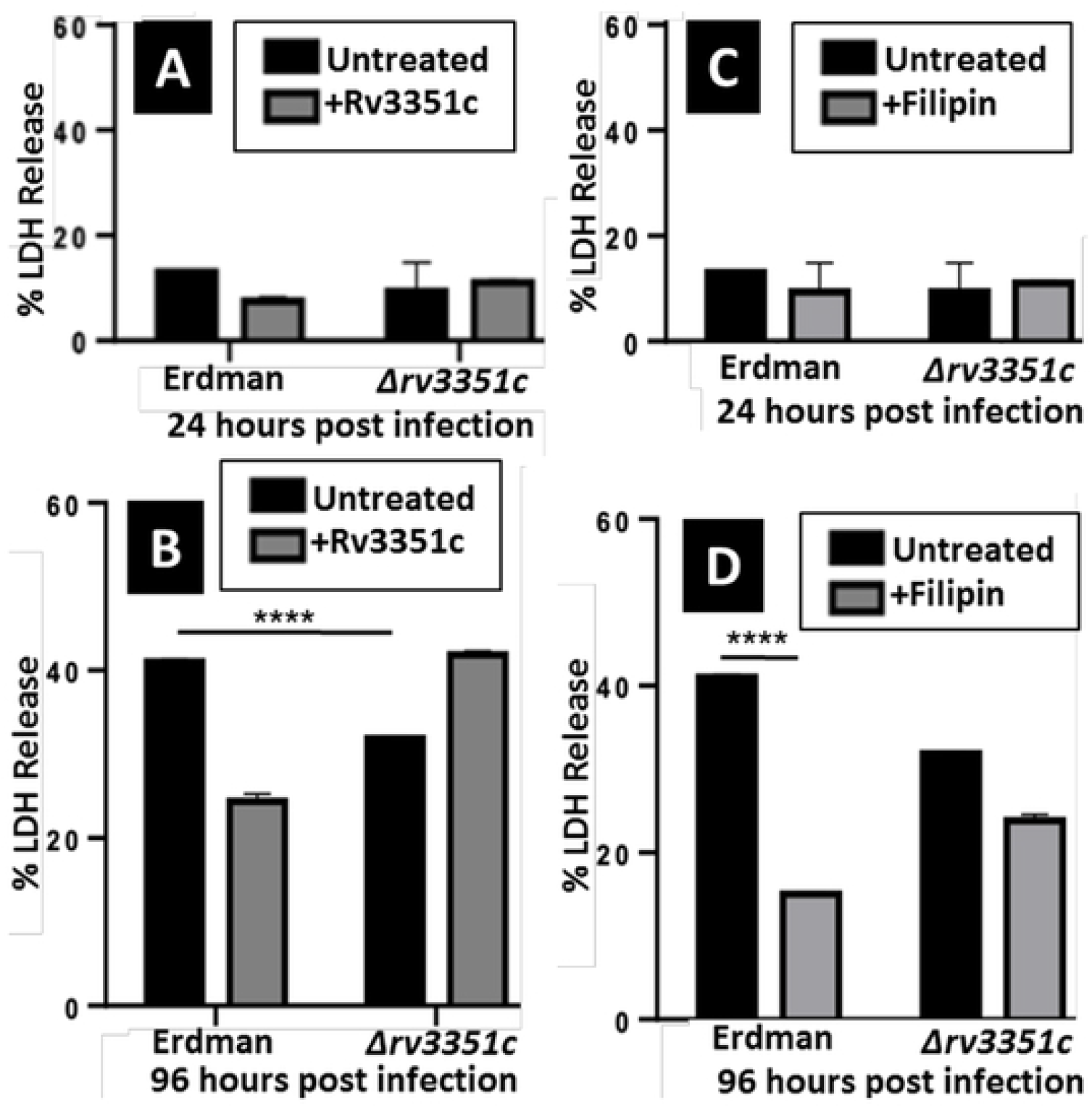
Addition of Rv3351c restores virulence of *Δrv3351c* in A549 cells. (A and C) LDH release 24 hpi from A549 cells infected with *M. tuberculosis* strain Erdman or *Δrv335l* treated with Rv3351c (A) or the cholesterol-binding agent, Filipin III (C). (B and D) LDH release 96 hpi, under the same conditions described for A and C, respectively. Data shown are the means of 3 experiments performed in triplicate, ****p-values: (p<0.001). Error bars represent one standard deviation.

### *Mycobacterium tuberculosis* Erdman and *Δrv3351c* bacilli attach to A549 type II pneumocytes via actin-mediated engulfment

To investigate if *rv3351c* influences *M. tuberculosis* entry into type II pneumocytes, we examined mycobacterial interactions at the surface of the host cell by transmission electron microscopy (TEM) and confocal microscopy. A549 cells were infected with *Δrv3351c* or the parent strain, *M. tuberculosis* Erdman. Images collected at 6 hpi suggest that both strains attach to philopodia and internalize through an actin-mediated mechanism (Figure 3, panels A and B). To test this hypothesis, A549 cells were allowed to attach to coverslips in 6-well dishes and were then infected with dsRed-expressing *M. tuberculosis* Erdman or GFP-expressing *Δrv3351c* bacilli at a MOI of 100. Preliminary experiments suggested that *M. tuberculosis* Erdman bacilli attached to the host cells and became internalized between 6 and 24 hpi. Specimens were fixed and stained with Alexa Fluor 488 or 546 phalloidin to visualize potential surface interactions via confocal microscopy. At 6 hpi, top virtual slices from z-stack images showed *M. tuberculosis* Erdman and *Δrv3351c* bacilli within coils of polymerized actin suggesting that both the mutant and parent strains interact with the alveolar epithelial host cell using a similar mechanism during initial infection (Figure 3, panels C and E).

**Figure 3.**
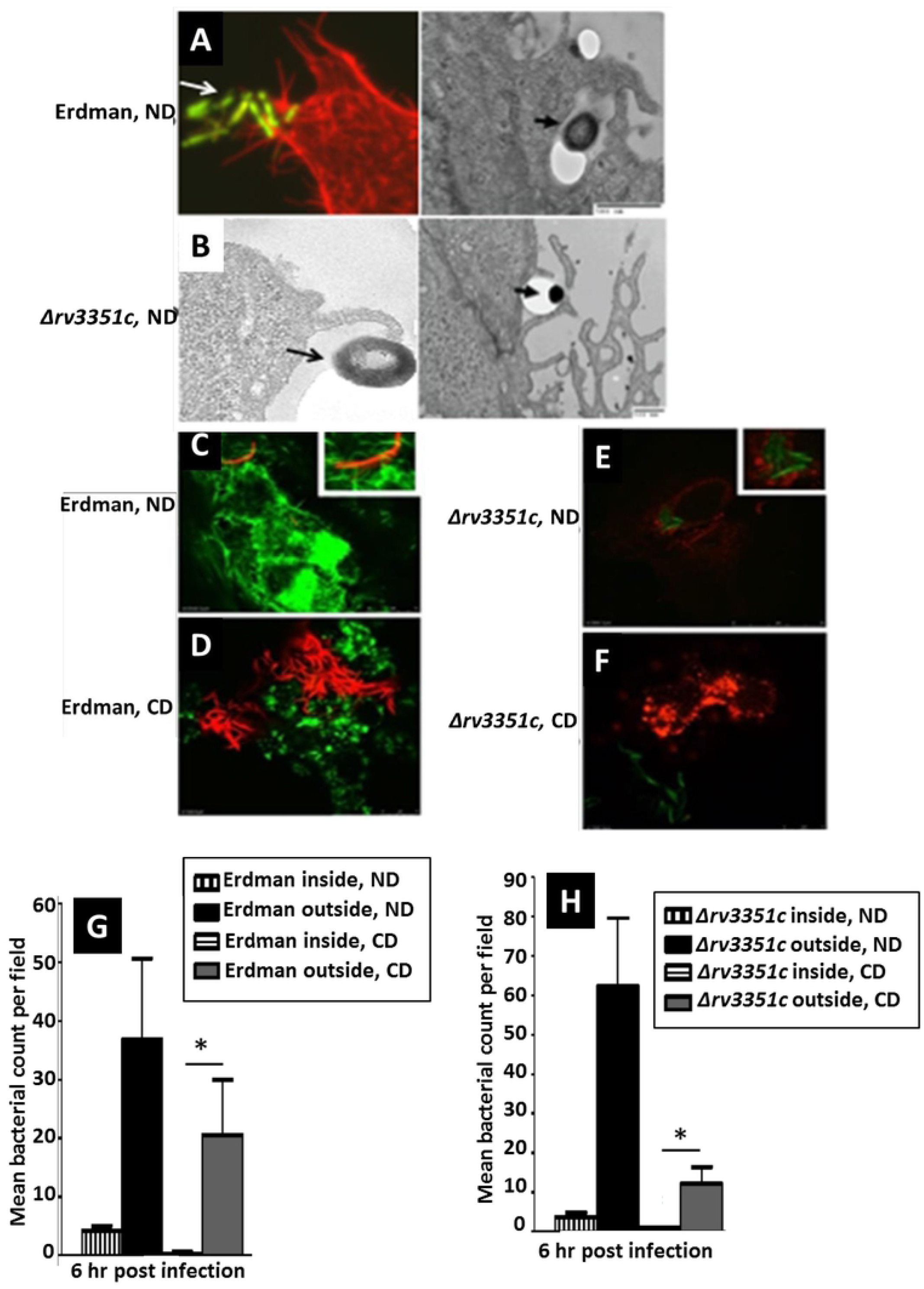
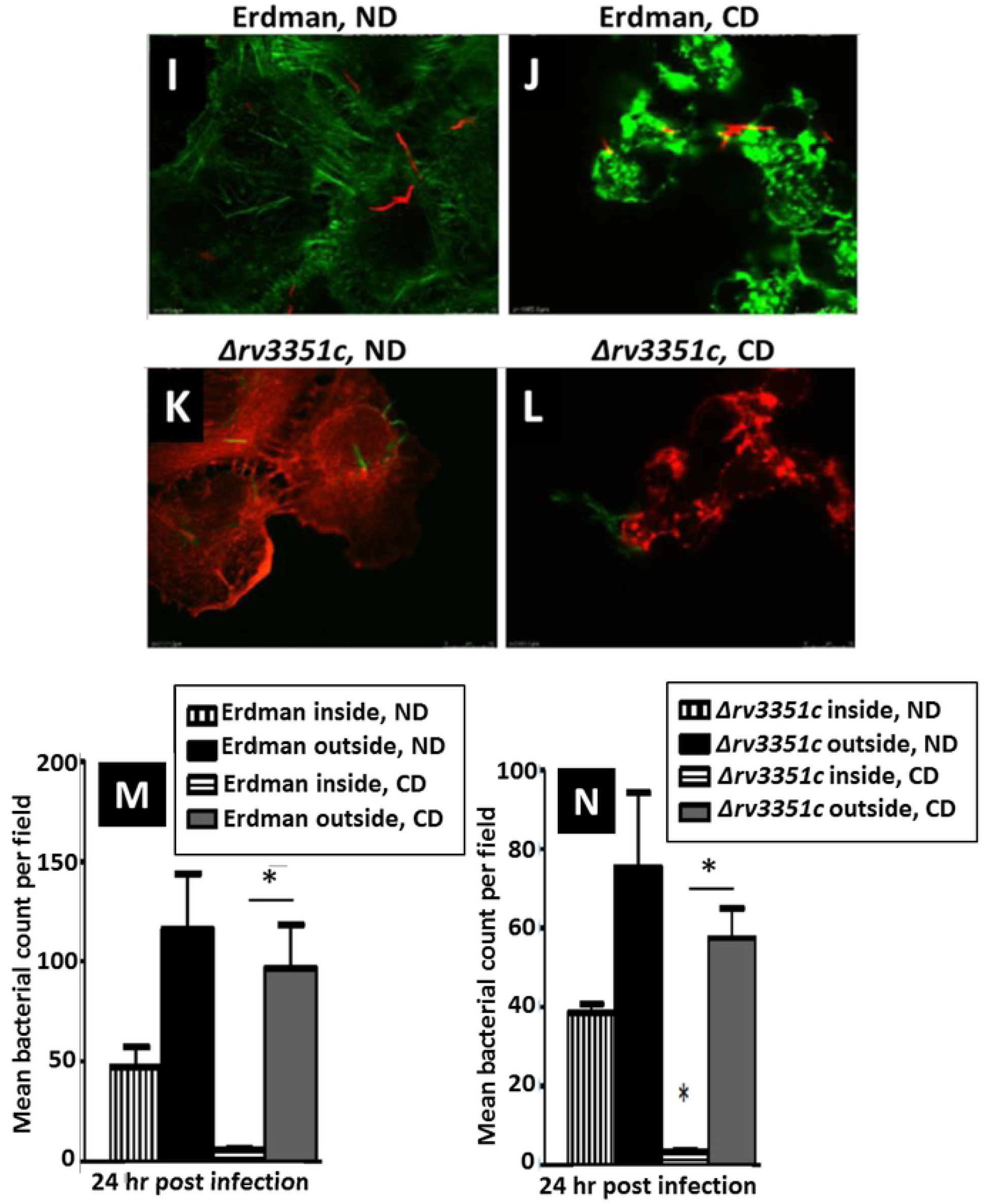
*Mycobacterium tuberculosis* bacilli attach to A549 cells via actin-mediated engulfment. A549 cells were infected at a MOI of 100 with *M. tuberculosis* Erdman or *Δrv3351c* bacilli and examined at the indicated times by confocal microscopy or TEM. (A) At 6 hpi, confocal z-stack imaging reveals phalloidin-stained red actin pseudopods wrapping around the *M. tuberculosis* Erdman bacilli (green) (left panel). TEM image showing *M. tuberculosis* Erdman bacilli interacting with host cell surface membrane projections (right panel, arrows indicate bacilli, magnification 7,000X). (B) At 6 hpi, TEM shows *Δrv3351c* bacilli interacting with host cell surface membrane projections (arrows indicate bacilli, magnification 10.000X [left figure] and 1,000X [right figure]). (C-F) At 6 hpi, confocal z-stack images reveal phalloidin-stained green (C) or red (E) actin pseudopods wrapping around red *M. tuberculosis* Erdman (C) and green *Δrv3351c* (E) bacilli at the surface of the untreated host cells (see insets on C and E). At 6 hpi, A549 cells treated with cytochalasin D show similar attachment numbers for both bacterial strains in the absence of actin polymerization (D and F). (G and H) Quantification from confocal studies comparing the same strains (parent or mutant) under the same conditions (cytochalasin D treated or untreated) shows no significant difference in numbers of bacteria adhering to host cells at 6 hpi, but no internalization of either strain, *p-values: (p<0.001). (I-L) At 24 hpi, z-stack images of cytochalasin - treated cells (J and L) show an absence of internalized *M. tuberculosis* Erdman (red) and *Δrv3351c* (green) bacilli while untreated A549 cells (I and K) have numerous internalized bacteria from both strains. (M and N) Quantification of the images in panels l-L revealed significant differences in numbers of internalized bacteria from cytochalasin D treated A549 cells compared to untreated cells infected with either strain, *p-values: (p<0.060). Error bars represent one standard deviation. Confocal image A was 100X magnification, C-F and l-L were captured at 63x magnification with a 3x zoom, and inset boxes on images C and E were 100X. ND, no drug added; CD, cytochalasin D added.

Previous observations have been made indicating actin-associated internalization of virulent *M. tuberculosis* strain H37Rv in A549 and mast cells (11,23,24). To determine if actin is critical for engulfment of both *M. tuberculosis* Erdman and *Δrv3351c*, the actin-depolymerizing agent cytochalasin D was utilized. Drug-treated A549 cells infected at a MOI of 100 with *M. tuberculosis* strains Erdman or *Δrv3351c* were examined at 6 and 24 hpi by confocal microscopy. These images revealed an absence of actin tendrils around either strain; however, no significant differences in the adherence of *M. tuberculosis* Erdman or *Δrv3351c* bacilli to the surface of host cells at either time point were observed with and without cytochalasin D suggesting that attachment can proceed in the absence of actin polymerization for both strains (Figure 3, panels D, F-N). At 6 and 24 hpi, the rates of internalization for *Δrv3351c* and *M. tuberculosis* Erdman bacilli were significantly reduced (essentially to baseline) compared to infected A549 cells not treated with cytochalasin D (p-value = 0.060 or <0.001, respectively; (Figure 3, panels I-N). Taken together, these data support that both *M. tuberculosis* Erdman and *Δrv3351c* bacilli utilize an actin-based mechanism for internalization, while attachment is not inhibited by the absence of polymerized actin for either strain.

### *Mycobacterium tuberculosis* Erdman and *Δrv3351c* bacilli co-localize with LAMP-2 and cathepsin-L in A549 cells at 12 to 96 hours post-infection

Thin layer chromatography overlay assays indicated that the Rv3351c protein can bind to hydrophobic components and glycosphingolipids in lipid extracts of A549 cells, consistent with the hypothesis that it binds lipid raft components. Previously, we reported that *Δrv3351c* does not survive as well as parent strain Erdman in type II pneumocytes (8). Combined with the results presented above, we hypothesize that Rv3351c induces lipid raft aggregation to enable *M. tuberculosis* Erdman bacilli to traffic advantageously in AECs. *Mycobacterium tuberculosis* bacilli inhibit phagosome-lysosome fusion in macrophages to escape killing. To test our hypothesis that *Mycobacterium tuberculosis* bacilli evade killing in epithelial cells by similar trafficking mechanisms, co-localization of Erdman and *Δrv3351c* bacilli with lysosomal-associated membrane protein 2 (LAMP-2) and the lysosomal protease cathepsin-L were examined in A549 cells.

Co-localization of GFP-expressing *M. tuberculosis* Erdman with LAMP-2 was examined by confocal microscopy from 6 to 96 hpi with some co-localization detected from 24 to 96 hpi (Figures 4, panels A-D; I-K). *Δrv3351c* bacilli were also detected co-localizing with LAMP-2 during the same time period (Figure 4 panels E-H; L-N). Co-localization of lysosomal markers with *M. tuberculosis* Erdman or *Δrv3351c* bacteria-containing compartments were quantified by immunoelectron microscopy using antibodies to cathepsin-L and LAMP-2. At 12 hpi, 60% of Erdman and 70% *Δrv3351c* bacilli in endosomes co-localized with cathepsin-L; however, at 72 hpi co-localization of Erdman dropped to 25%, while the mutant dropped only to 45% (Figure 5, panels A-E). Endosomes containing strain Erdman co-localized with LAMP-2 much less frequently than those containing *Δrv3351c* bacteria. Co-localization was 3% for Erdman and 15% for *Δrv3351c* at 12 hpi, while at 72 hpi the percentages increased to 13% for Erdman, but to 48% for the mutant (Figure 5, panel F).

**Figure 4.**
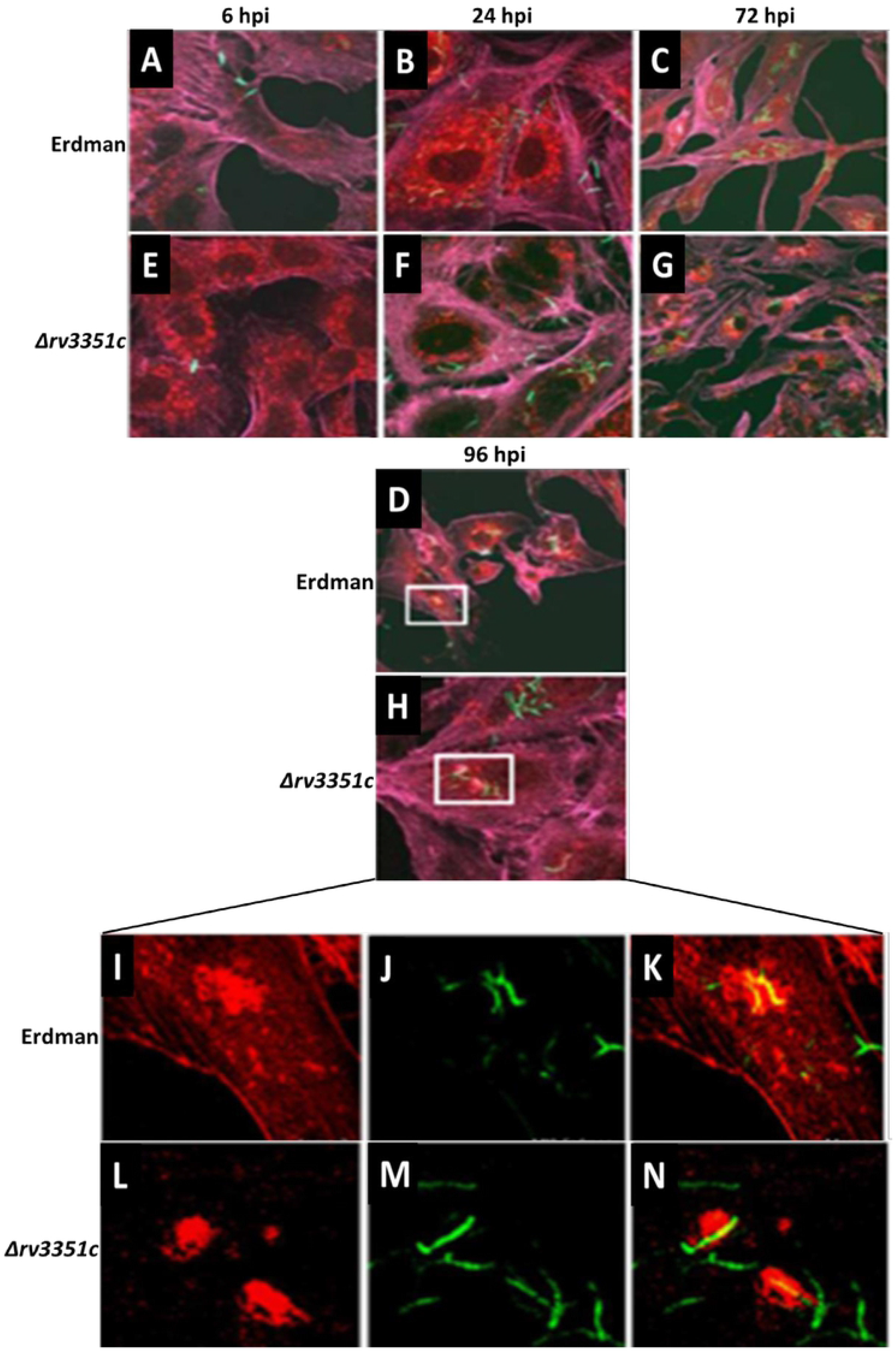
Confocal microscopy demonstrates intracellular differences between *M. tuberculosis* Erdman and *Δrv3351c* bacilli in A549 cells. (A-H) At 6, 24, 72 and 96 hours post infection, A549 cells were prepared for confocal microscopy as described. *M. tuberculosis* Erdman (A-D) and *Δrv3351c* (E-H) (green) were examined following staining with LAMP-2 (red) for co-localization (yellow). Boxed regions in panels D and H identify the regions displayed in overlay series (l-K; Erdman and L-N; *Δrv3351c*). Confocal images A-H were captured at 63x magnification while images l-N were 100X magnification.

**Figure 5.**
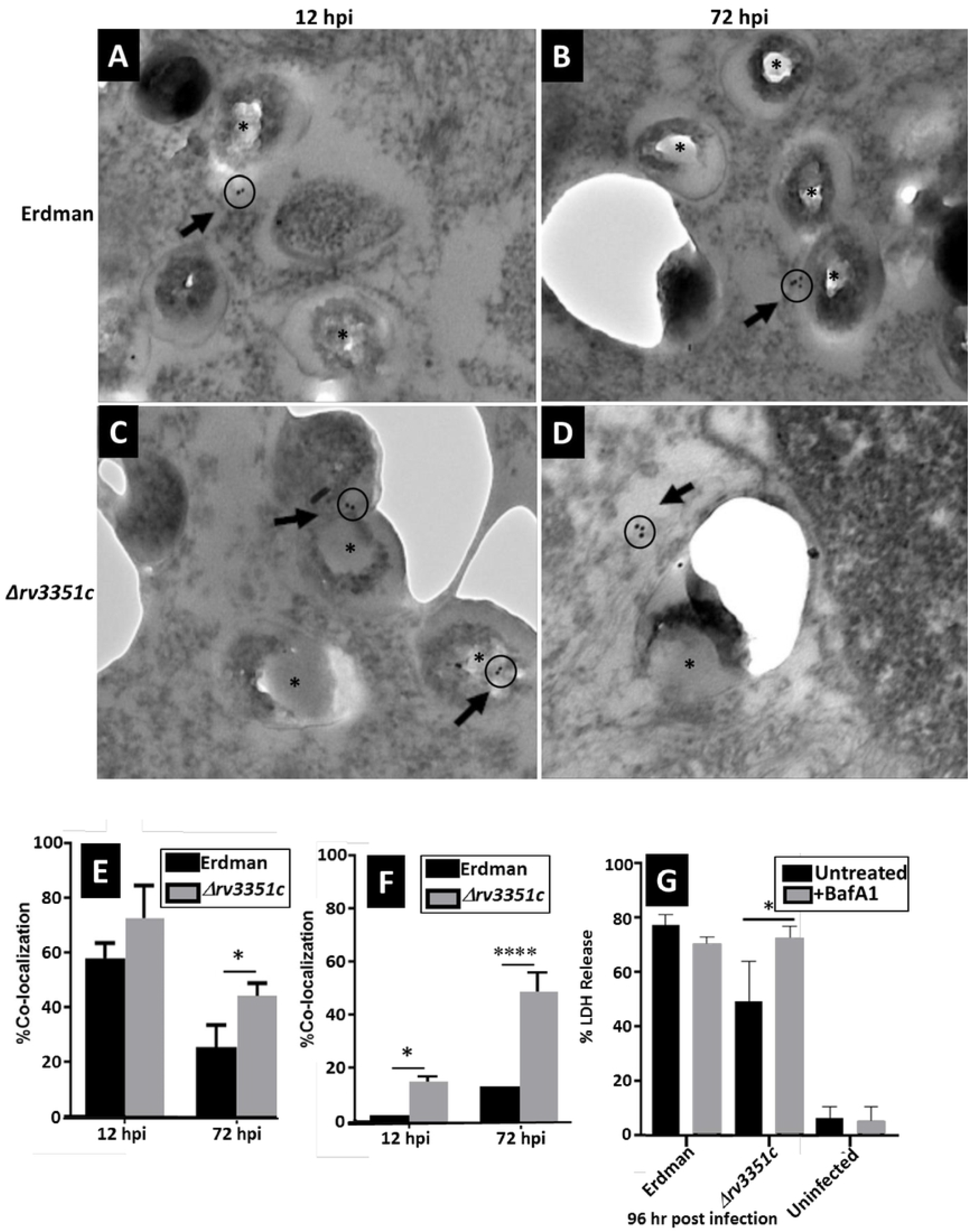
Immuno-electron microscopic quantification of mycobacterial co-localization with cathepsin-L or LAMP-2. Following A549 infection with either *M. tuberculosis* strain Erdman or *Δrv3351c*, specimens were prepared for IEM. Cathepsin-L was labeled with 12-nm gold particles. Shown are representative images (A-D). Bacilli are denoted by (*) and gold particles are identified with an arrow in each panel. Percentages of bacteria co-localized with either cathepsin-L (E) or LAMP-2 (F) were quantified, ****p-values: (p<0.0001), *p-values: (p<0.012); scale bars = 500 nm for panels A and B or 100 nm for panels C and D). (G) Inhibiting lysosomal acidification with Bafilomycin A1 (100 nm) restored the *Δrv3351c* LDH phenotype to wild type by 72 hpi, *p-values: (p<0.05). Error bars represent one standard deviation. TEM images are shown at a magnification of 7,000X.

To assess whether the previously-observed reduced cytopathy of the *Δrv3351c* infected cells correlates with increased killing of the mutant by the acidic lysosomal environment, the vacuolar type H+-ATPase inhibitor Bafilomycin A1 was added to A549 monolayers prior to infection. Inhibiting lysosomal acidification with Bafilomycin A1 increased released lactate dehydrogenase activity from cells infected with *Δrv3351c* bacilli to the level detected in cells infected with Erdman, but had no effect on cells infected with Erdman bacilli at 72 hpi (Figure 5, panel G). Increased cytopathic effects of the mutant in cells treated with Bafilomycin A1 is consistent with greater survival of the bacteria within phagolysosomes. Studies showed no deleterious effects produced by the drug alone at the concentration used (100 nm) in uninfected cells (data not shown).

### *Mycobacterium tuberculosis* Erdman and *Δrv3351c* bacilli induce production of autophagy proteins in A549 cells

We previously showed that induction of the autophagy pathway is crucial for *M. tuberculosis* Erdman survival in A549 cells. To determine if Rv3351c was directly involved in autophagy induction, A549 cells were infected with strain Erdman or *Δrv3351c* bacilli expressing dsRed or GFP, respectively for 12 (Figure 6A and C) or 24 hours (Figure 6B and D). Confocal microscopic examination revealed greater amounts of LC3 puncta at 12 hpi in *Δrv3351c*-infected cells than those infected with Erdman, but the opposite trend was observed at 24 hpi (Figures 6E and F, respectively).

**Figure 6.**
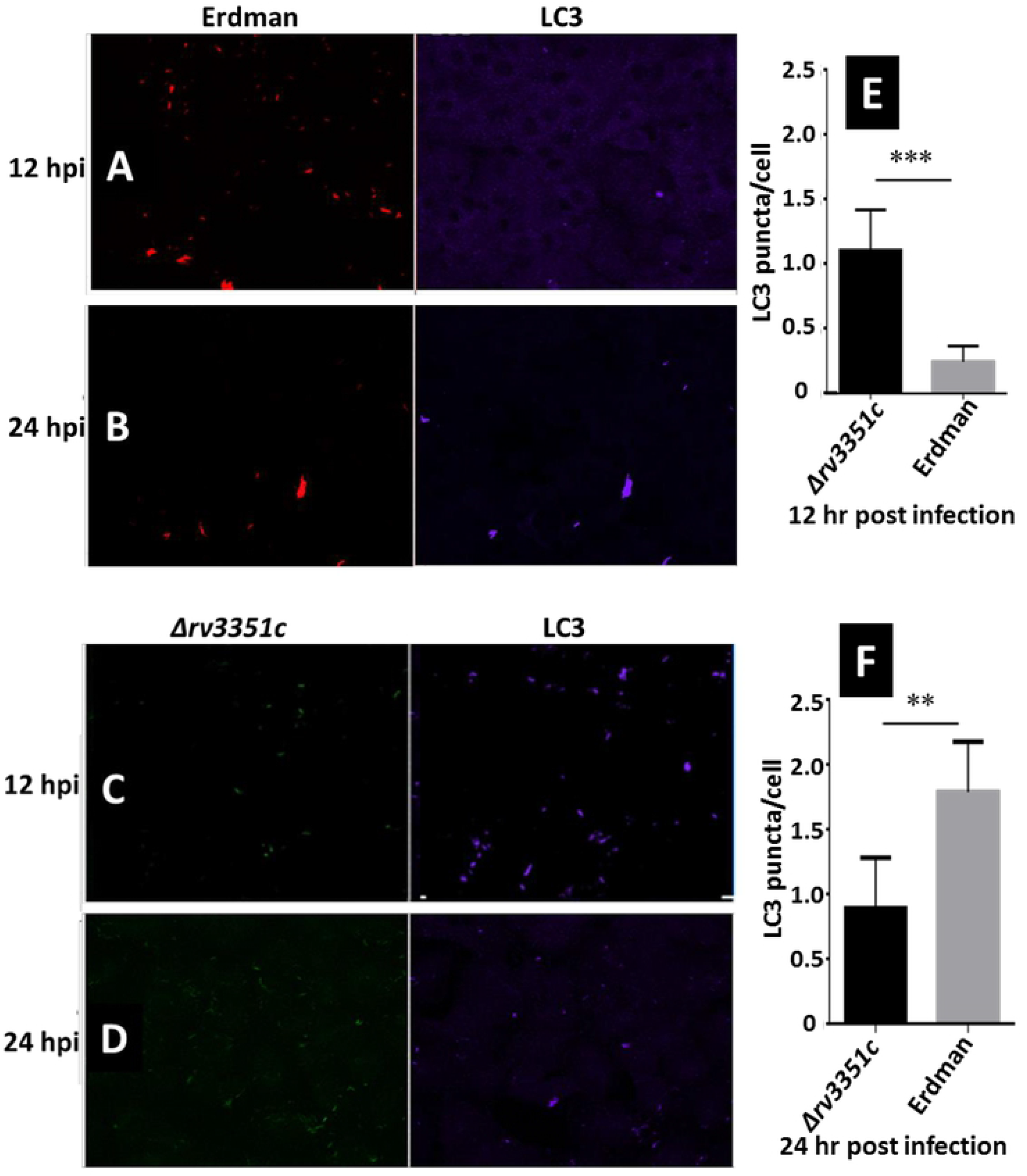
Confocal microscopy images demonstrate *M. tuberculosis* Erdman and *Δrv3351c* bacilli induce autophagy marker LC3 in A549 cells. (A-D) A549 cells were infected with strain Erdman or *Δrv3351c* expressing DsRed or GFP, respectively for 12 (A and C) or 24 hours (B and D). LC3 immuno-labeling appears purple. DsRed-expressing *M. tuberculosis* bacilli appear red. GFP-expressing *Δrv3351c* bacilli appear green. Punctate LC3 staining is observed in cells infected with strain Erdman (A and B) and *Δrv3351c* (C and D). (E and F) Puncta of LC3 were quantified at 12 and 24 hpi. Images are representative of three experiments with similar results Data shown are the means of three experiments performed in triplicate, **’p-values: (p<0.005), **p-values: (p<0.05). Error bars represent one standard deviation. Confocal images were captured at 40X magnification.

To examine conversion of LC3 forms, A549 cells were incubated for 12 or 24 hours with *M. tuberculosis* Erdman or *Δrv3351c* at a MOI of 100. After extracellular bacteria were removed, Autophagy levels were analyzed by LC3 immunoblotting assay with GAPDH as a loading control (Figure 7A and B). The LC3-II/GAPDH ratio is higher after *Δrv3351c* or Erdman infection than uninfected controls at 12 hpi (Figure 7A and C); however, the LC3-II/GAPDH ratio is higher at 12 hpi for A549 cells infected with *Δrv3351c* than Erdman or cells induced for autophagy by amino acid starvation with EBSS (Figures 7A and C). This difference disappears by 24 hours (Figures 7B and D). Sets of cells were treated for two hours with Bafilomycin A1 after infection to examine the effects of inhibiting autophagic flux. Addition of the drug did not significantly impact LC3-II conversion levels (Figure 7).

**Figure 7.**
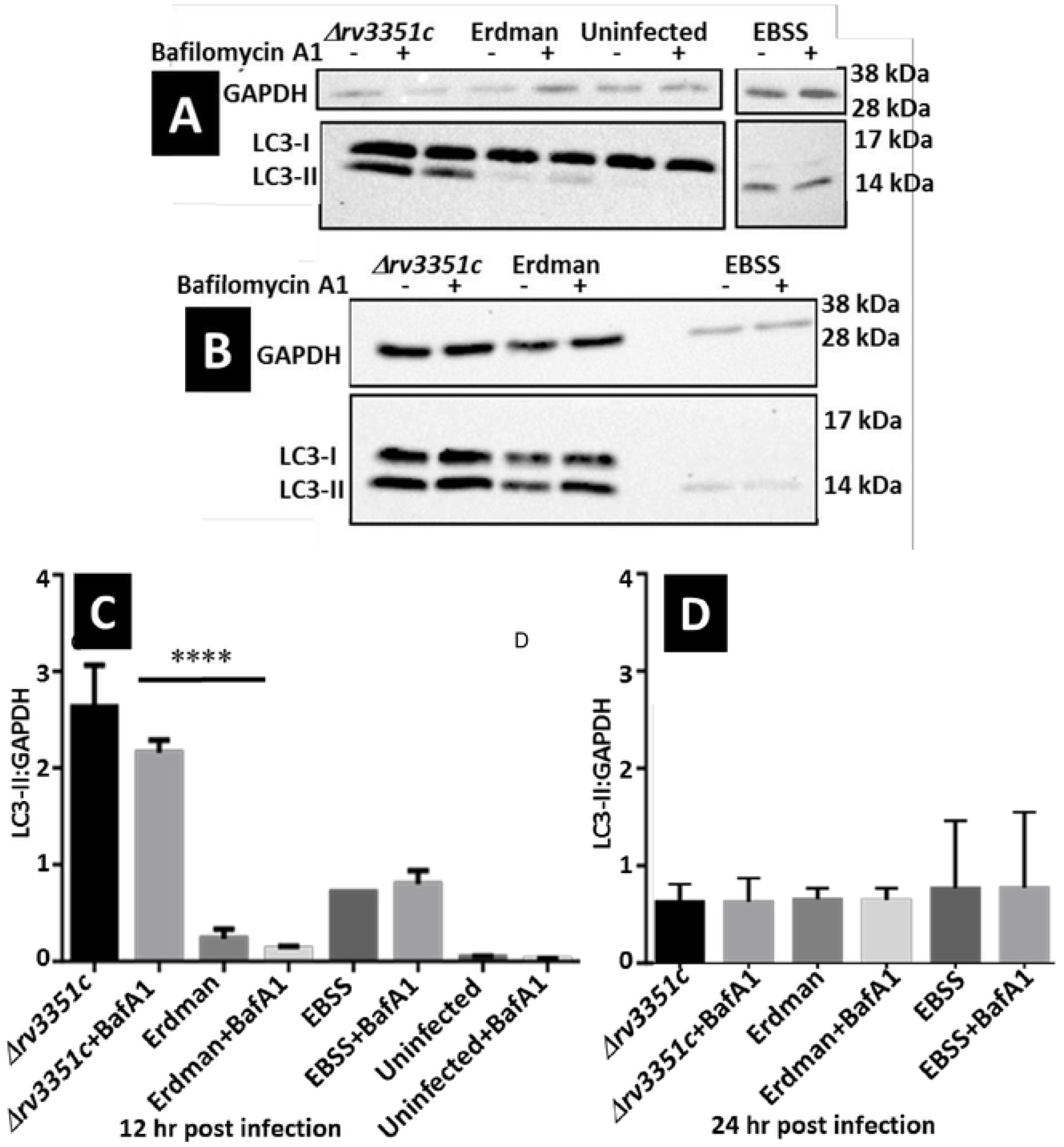
Western-blot analysis demonstrates LC3-II conversion. (A and B) Cell lysates were obtained from infected A549 cells, uninfected cells, or cells under starvation (EBSS) conditions at 12 and 24 hours in the presence or absence of autophagy inhibitor Bafilomycin A1 (100 nM) as described. (C and D) LC3-II to GAPDH loading control ratios were quantified. Blots are representative of two experiments with similar results. Data shown are the means of two experiments, ****p-values: (p<0.0001). Error bars represent one standard deviation.

Due to the early observed differences in LC3-II production between A549 cells infected with *Δrv3351c* versus Erdman, we sought to determine if there were differences in the production of other autophagy proteins. ULK1 is involved in autophagosome formation upstream of LC3 recruitment (25). Densitometric analysis shows differences in the ULK1/GAPDH ratios in Erdman and uninfected cells at 24 hpi, while no differences are seen between *Δrv3351c* and uninfected cells at that time point (Figure 8B and D), indicating *Δrv3351c* may not be trafficking through the autophagy pathway.

**Figure 8.**
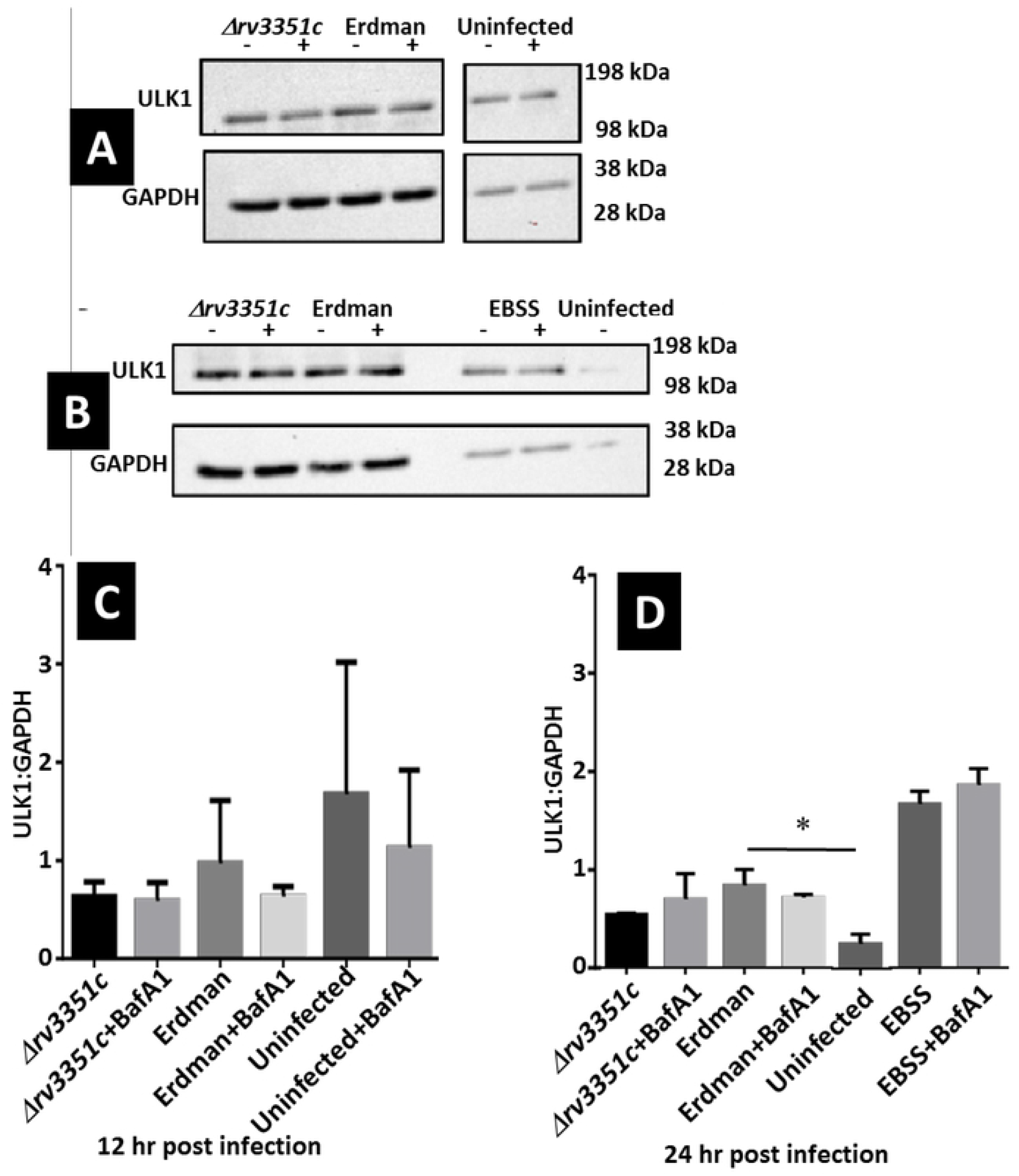
Western-blot analysis demonstrates induction of ULK1. (A and B) Cell lysates were obtained from infected A549 cells, uninfected cells, or cells under starvation (EBSS) conditions at 12 and 24 hours in the presence or absence of autophagy inhibitor Bafilomycin A1 (100 nM) as described. (C and D) ULK1 to GAPDH loading control ratios were quantified. Data shown are the means of two experiments, *p values: (p< 0.05). Error bars represent one standard deviation.

### Cytokine profiles of A549 cells infected with *M. tuberculosis Δrv3351c* or Erdman bacilli

In response to bacterial infection, epithelial cells at the mucosal surface secrete cytokines important in lung defense and inflammatory responses (26). Tumor necrosis factor alpha (TNF-α) plays a critical role in the control of *M. tuberculosis* (26); however, these bacteria can also inhibit host cell TNF-α production evade anti-tuberculosis immunity (27). In response to *M. tuberculosis* infection, alveolar epithelial cells secrete IL-8 and MCP-1 to recruit reinforcing immune-system cells (28-30). Cytokine production in A549 cells was examined following infection with *M. tuberculosis* Erdman or *Δrv3351c* at a MOI of 100. At 12, 24, and 72 hpi, supernatants were assayed for TNF-α, IL-8, and MCP-1 production. At 72 hpi, Erdman or *Δrv3351c* infected cells produced similar high levels of MCP-1 and IL-8 (Figure 9B and C). The *Δrv3351c* bacilli stimulated significantly-higher levels of TNF-α compared to infection with strain Erdman (Figure 9A). Induction of TNF-α by *Δrv3351c* infection was blocked by addition of Rv3351c protein (Figure 9A). Interestingly, infection with nontuberculous mycobacterium species, *M. smegmatis* also induced high levels of TNF-α (Figure 9A). This species does not possess a close *rv3351c* homolog.

**Figure 9.**
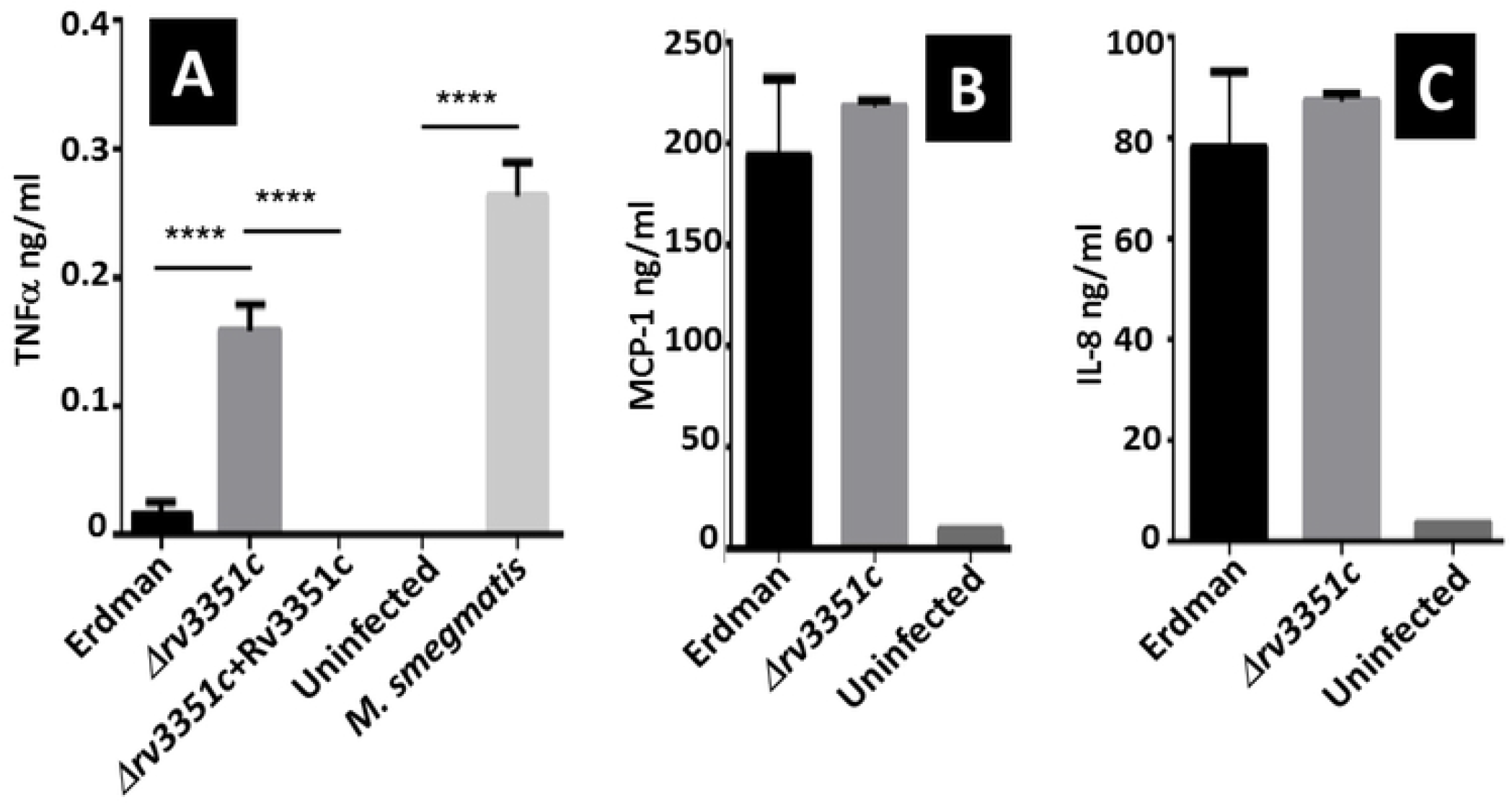
ELISA of infected A549 supernatants. (A-C) A549 cells produce TNF-α, MCP-1, and IL-8 in response to *M. tuberculosis* Erdman or *Δrv3351c* 72 hpi. (A) *M. tuberculosis* Erdman induced low production of TNF-α while *Δrv3351c* and the non-pathogenic species *M. smegmatis* generated a high TNF-α responses. The addition of 5 μg Rv3351c protein to *Δrv3351c* reduced expression of TNF-α to wildtype Erdman levels. MCP-1 (B) and IL-8 (C) are induced to comparable levels by infection with Erdman or *Δrv3351c*. Data shown are the means of three experiments, ****p-values (p<0.0001). Error bars represent one standard deviation.

### Rv3351c is detected in the outer membrane and recognized by the host immune response in human tuberculosis patients

Due to its role in the aggregation of epithelial cell lipid rafts, Rv3351c likely localizes to the cell surface or is extracellularly secreted by *M. tuberculosis*. Previous studies have shown mycobacterial surface protein HBHA interacts with alveolar epithelial cells and induces humoral immune responses in humans and elicits a protective immune response in mice (31,32). Probing *M. tuberculosis* Erdman cell lysate fractions with an antibody raised against recombinant Rv3351c protein detected a prominent band of the expected size for Rv351c in the membrane fraction, and a faint band in the cytoplasmic fraction, but not in the culture supernatant (Supplemental Figure 2). If Rv3351c is surface-associated and antigenic, the host may develop antibodies against the protein. Sera from humans with culture-confirmed active tuberculosis, latent tuberculosis, or negative controls were screened by ELISA. Latent tuberculosis infection is defined as a state of persistent immune response to stimulation by *M. tuberculosis* antigens without evidence of clinically-manifested active tuberculosis (33). Patients with likely latent infection had negative bacteriological tests and the presumptive diagnosis is based on a positive result of a tuberculin skin test or interferon-gamma release assay indicating an immune response to *M. tuberculosis* infection (33). Results indicate that human sera from patients with active, but not latent tuberculosis, produced antibodies against the Rv3351c protein (Figure 10).

**Figure 10.**
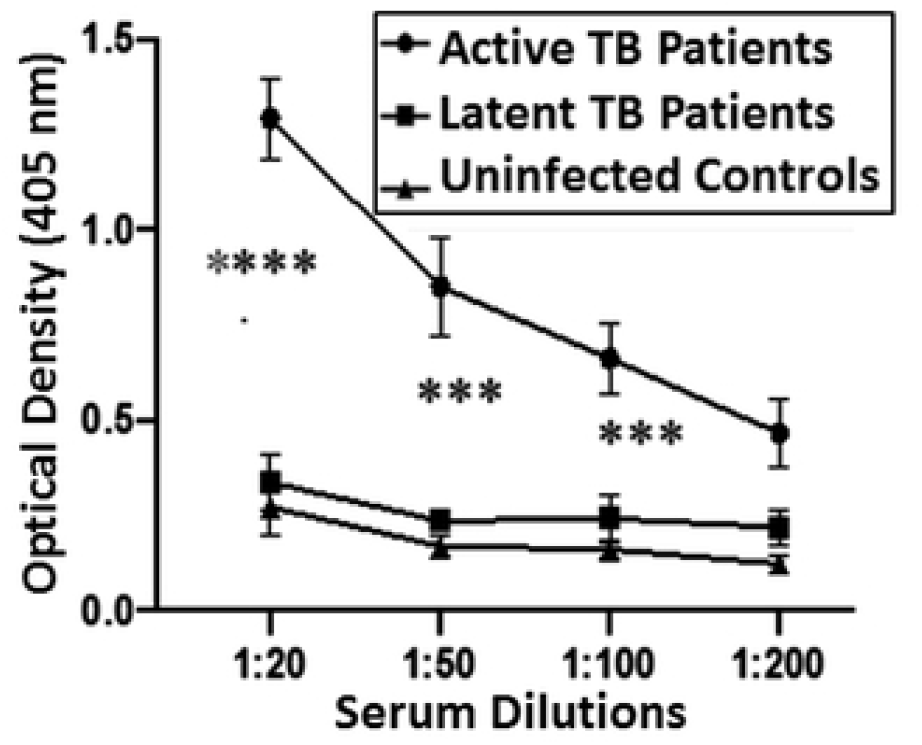
Screening of human sera from patients with culture-confirmed active versus latent TB and negative control volunteers by ELISA show that these patients produce antibodies against the Rv3351c protein. Images are representative of three experiments with similar results, ***’p-values: (p<0.0001), ***p-values: (p<0.001). Error bars represent one standard deviation.

### *Δrv3351c* bacilli are less virulent in the mouse aerosol-infection model

It was observed that BALB/c mice intratracheally-infected with the *Δrv3351c* bacilli have fewer replicating bacilli in lungs and fewer bacilli disseminated to the spleens and particularly the livers by 21 days post-infection compared to control infections with the complemented mutant strain, *CMΔrv3351c* (Figure 11). Infection with either the mutant or complemented strain generated similar immune cell infiltration by the three-week post-infection time point (Supplemental Figure 3).

**Figure 11.**
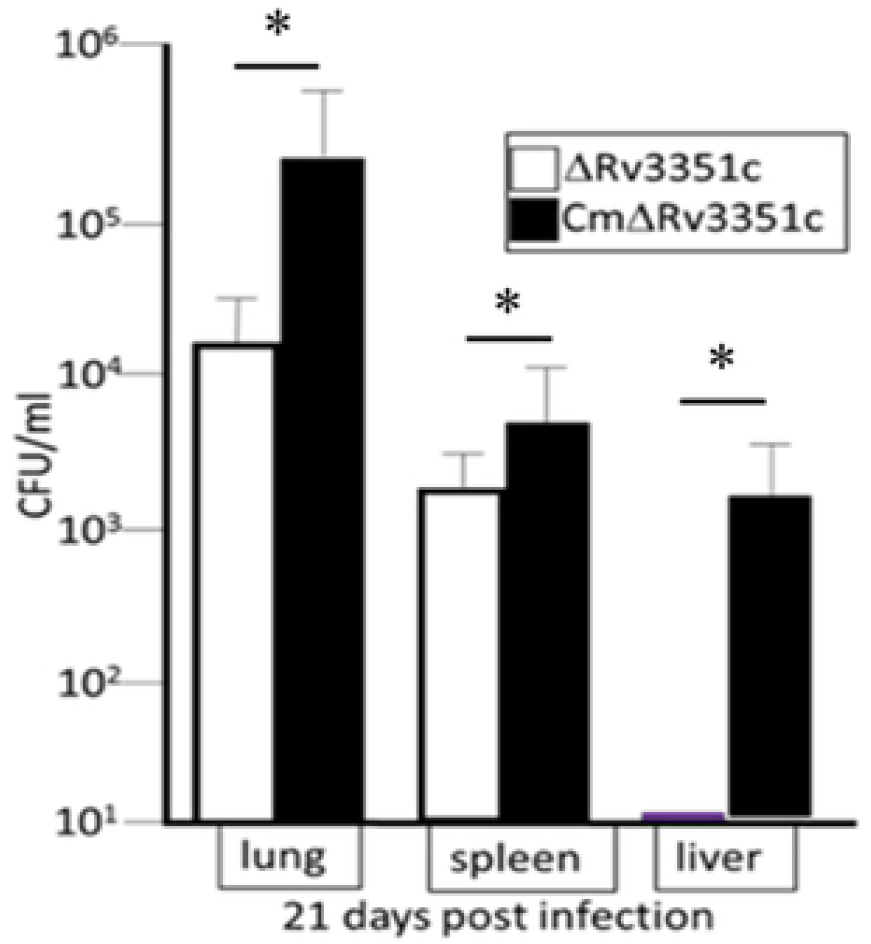
BALB/c mice intranasally-infected with the *Δrv3351c* bacilli have fewer replicating bacilli in lungs and fewer disseminated to the spleen and liver 21 days post-infection compared to control infections with the complemented mutant, CM¿rv3351c. The intratracheal dose of these strains given was 1X10^5^ CFU in 0.025 ml PBS. Tissues were collected after euthanasia and processed for viable counts as described, *p values (p< 0.05). Error bars represent one standard deviation.

## Discussion

To better characterize the role of type II pneumocytes in tuberculosis, particularly during early infection, detailed studies of the interaction between the bacilli and alveolar pneumocytes at the cellular level are needed. In the experiments described herein, a process of bacterial attachment, internalization and trafficking through type II pneumocytes has been established, including measurements of bacterial and host cell viability. Using a deletion mutant of *M. tuberculosis* Erdman lacking the *rv3351c* gene (*Δrv3351c*), we previously determined that replication in alveolar epithelial cells as well as macrophages is decreased compared with wild type or complemented control strains (9). In the present study, the attachment, internalization and trafficking within alveolar epithelial cells by *Δrv3351c* bacilli were compared to those of its parental strain. In addition, the mutant strain was shown to disseminate less efficiently from the lungs of infected mice compared to its complement, and the Rv3351c protein induced cellular and humoral immune responses during infection.

### rv3351c involved in lipid raft faggregation and intracellular trafficking of *M. tuberculosis*

We previously demonstrated microscopically that viable *M. tuberculosis* Erdman bacilli induce lipid raft aggregation on infected A549 cells to similar or greater levels than lipid raft super aggregator LLO (9). Additionally, we showed that culture filtrates from infected A549 cells also induce lipid raft aggregation when added to fresh monolayers indicating the responsible factor is mycobacterial and likely secreted during or prior to infection (9). In this current study, we demonstrate that infection with *Δrv3351c* cells did not as efficiently aggregate lipid rafts compared to the parent strain, but recombinant Rv3351c protein alone induced similar levels of lipid raft aggregation on A549 cell plasma membranes as the LLO positive control when added at equal concentrations (Figure 1). Interestingly, the difference in lipid raft aggregation between A549 cells infected with the wild type and *Δrv3351c* bacilli did not appear at the gross level to affect the method or rate of bacterial attachment and internalization, but did alter the intracellular trafficking pattern between the two strains. Thus, the overall studies described here provide for a more thorough understanding of the role Rv3351c in the process of *M. tuberculosis* attachment, internalization and trafficking within type II pneumocytes. However, these data also contribute to the body of knowledge indicating a larger role for alveolar pneumocytes; they may not simply provide a barrier to infection, but may also contribute to the pathology associated with tuberculosis (34-37). What was surprising is the apparent role this protein plays in modifying the host cell membrane to induce lipid raft aggregation, and thus plays a major role in the optimal trafficking that is not observed with the mutant strain; potentially causing the mutant bacteria to attach to, enter the pneumocyte, and traffic through a sub-optimal pathway that is sub-optimal for bacterial survival and spread.

The process of attachment and internalization is readily observed with both *M. tuberculosis* Erdman and *Δrv3351c* bacilli between 6 and 24 hpi. Other studies have shown mycobacterial internalization in these cells occurring as early as 2 hpi (11,38). Thus, our results are comparable, although different *M. tuberculosis* strains and different methods of quantification make direct comparisons difficult. There has been some discussion regarding the actual mechanism of internalization in type II pneumocytes with studies suggesting a membrane ruffling process may occur (23). In this study, microscopic examination did not reveal evidence of ruffling but did support previous observations of actin-pseudopod engulfment (23). Furthermore, addition of the actin-depolymerizing agent, cytochalasin D, resulted in a significant reduction in internalized mutant and wild-type bacteria at 24 hpi (Figure 3). This suggests that *M. tuberculosis* Erdman and *Δrv3351c* bacilli internalize using the same actin-mediated mechanism.

Although internalization was reduced by the addition of cytochalasin D, adherence of mycobacteria to treated host cells was not significantly different from the infected, untreated host cells. Not surprisingly, these data suggest that *M. tuberculosis* bacilli can attach to alveolar epithelial cells through multiple mechanisms, with actin-pseudopod engulfment being the preferred and perhaps more efficient means of intimate cellular association/internalization. The timeline for the process of attachment and internalization in human macrophages has been reported to occur within one hour (39). The longer time period required for bacterial internalization in the pneumocyte compared to the macrophage is not unexpected given that epithelial cells are not phagocytic by nature and thus an alternative, potentially slower means of internalization is at work. The precise nature of the attachment and internalization mechanism is currently being investigated.

Similar rates of attachment and internalization by Erdman and *Δrv3351c* bacilli were initially observed, however, analysis of the trafficking pathway demonstrated that the mutant bacilli are more often associated with endosomes possessing lysosomal markers LAMP-2 and cathepsin-L compared to the parent strain at 12 and 72 hpi (Figure 5). Similarly, inhibiting lysosomal fusion restores the mutant phenotype to wild type levels. Thus, the mutant bacilli may be exposed to higher levels of degradative lysosomal enzymes that are inhibitory or ultimately lethal, helping to explain the previously-observed decrease in intracellular mutant bacterial numbers observed over time (8). The *rv3351c* gene product might facilitate efficient trafficking by providing exposure to the appropriate host cell ligand associated with lipid rafts. Early association of mutant bacteria with LC3, and their ultimate delivery to the lysosome could suggest a mechanism similar to LC3-associated phagocytosis (LAP), in which LC3 and other autophagy proteins target single membrane bacteria-containing compartments to the lysosome for degradation (40). *Mycobacterium tuberculosis* and the related fish pathogen *M. marinum* are both targeted by LAP in macrophages, and mutants of *M. tuberculosis* bacilli lacking *cpsA* are degraded by LAP, while wild type bacilli are able to subvert this process (41,42). *Legionella dumoffii, Shigella flexneri*, and *Yersinia pseudotuberculosis* are all intracellular pathogenic bacteria that are targeted by LAP in non-phagocytic cells (43). ULK1 is a protein specific to the autophagy pathway (44), and is not essential in the LAP pathway. The inability of *Δrv3351c* to induce ULK1 above uninfected cell levels supports the hypothesis that *Δrv3351c* is trafficked through the LAP pathway (Figure 8D). While we have previously shown that *M. tuberculosis* Erdman bacteria are contained in double-membrane autophagosomal compartments, further investigation into the structure of single-membrane *Δrv3351c*-containing compartments needs to be completed, as well as examining bacterial association with other markers specific to either LAP or canonical autophagy.

### Rv3351c manipulates cellular and humoral immune responses and promotes *M. tuberculosis* extrapulmonary dissemination

Studies with other intracellular bacterial pathogens have shown that initial target cell membrane lipid raft aggregation may be a means to manipulate the host immune response; for example, expression of pro-inflammatory cytokines such as TNF-α were significantly decreased after infection of murine macrophages and human intestinal epithelial cells with *Brucella* sp. and *Campylobacter jejuni*, respectively (45,46). We have confirmed these observations and observed that A549 cells infected with *M. tuberculosis* Erdman induce TNF-α production to a significantly lower level than when infected with *Δrv3351c* (Figure 9), but production was again significantly reduced when Rv3351c was added to *Δrv3351c* during the infection. These data suggest the aggregation of lipid rafts induced by Rv3351c provide an additional mechanism by which *M. tuberculosis* modulates the host cellular immune response.

HBHA was the first pneumocyte-targeting *M. tuberculosis* protein to be identified and shown to be involved in mycobacterial attachment and intracellular trafficking. However, due to its surface localization and likely constitutive expression, this protein also was shown to induce humoral immune responses in human TB patients and subsequently to be an effective mucosal vaccine target (31,32). HBHA-deficient mutant strains of *M. tuberculosis* also replicate to lower numbers in the lungs of aerosol-infected mice compared with a wild-type strain and disseminate from the lungs less efficiently (13). Similarly, in this study, we showed that in addition to its role in pneumocyte internalization and trafficking, Rv3351c induces a humoral response in human TB patients (Figure 10), and the *Δrv3351c* mutant strain replicates to lower numbers in the lungs of mice and disseminates less effectively. Thus, these data infer that, like HBHA, Rv3351c functions extracellularly and is eventually targeted by the host immune system during infection.

Thus, with HBHA, our studies of Rv3351c support a hypothesis that *M. tuberculosis* has mechanisms to target the alveolar epithelium, enhancing the importance of these cells in the disease process. Elucidation of this process continues to be explored *in vitro* and *in vivo*.

## Materials and methods

### Ethics statement

These studies were conducted in accordance with the recommendations in the Guide for the Care and Use of Laboratory Animals and the American Veterinary Medical Association (AVMA). All animal experiments were performed with the approval of the Institutional Animal Care and Use Committee at the University of Georgia (NIH Animal Welfare Assurance Number: A3437-01).

### Bacterial strains

*Mycobacterium tuberculosis* strain Erdman was obtained from the Tuberculosis/Mycobacteriology Branch of the Centers for Disease Control and Prevention. The *M. tuberculosis* strains were grown in Middlebrook 7H9 broth supplemented with 0.5% glycerol, 0.05% Tween 80, 0.5% bovine serum albumin (fraction V, Boehringer-Mannheim), and 0.085% NaCl. Strain *Δrv3351c* was developed from strain Erdman using the method of Braunstein *et al*. (47) and is described in Pavlicek *et al*. (8). The *Δrv3351c*-complementing strain (*CMΔrv3351c*) was constructed by amplifying the *rv3351c* gene plus 366 bp of upstream and 14 bp of downstream DNA with primers 5’-GTG AAG CTT ATA CTG GTG AAG GTT TGCGC-3’ and 5’-GGA ATT CTT CGA CTG CTG GCG GAG-3’ and inserting the amplicon into the integrative plasmid pMV306 (48). This plasmid was then electroporated into strain Erdman and transformants were selected for by inclusion of kanamycin (25 μg/ml) in the medium (48). All virulence assessments were comparable between the parent and complemented strains on infected AECs and macrophages (8). For confocal microscopy, *M. tuberculosis* strains Erdman and *Δrv3351c* were transformed with replicating plasmid pFJS8gfpmut2 expressing green fluorescent protein (49) or replicating plasmid pGCRed2 expressing dsRed2, and maintained by inclusion of 50 μg/ml kanamycin or 50 μg/ml hygromycin, respectively. Plasmid pGCRed2 was a generous gift from Dr. Mary Hondalus, Department of Infectious Diseases, University of Georgia. Bacterial plating studies utilized Middlebrook 7H11 agar supplemented with 0.5% glycerol, 0.05% Tween 80 and 1 x ADS (8).

### Epithelial cell infection

A549 human type II alveolar epithelial cells were obtained from ATCC (CCL-185) and maintained in DMEM supplemented with 5% FBS at 37°C in 5% CO_2_. Twenty-four hours prior to infection, A549 cells were seeded into 6-well Costar^®^ dishes at 1.0 x 10^6^ cells per well. On the day of infection, epithelial cell monolayers were first prepared for a cold-synchronized infection by incubating at 4°C for one hour. Cells were then infected at a MOI of 100 (100 bacteria per host cell) with *M. tuberculosis* strains. To disperse the inoculum, bacteria were passed through an insulin syringe directly into the appropriate wells. After a one-hour incubation at 4°C post infection, cells were incubated at 37°C in 5% CO_2_; this was considered time point 0. Infections were maintained for up to 96 hours with samples examined at the indicated times. For actin-depolymerization experiments, cytochalasin D (1 μM final) was added to cells two hours prior to cold-synchronization and infection at a MOI of 100 and maintained in culture medium throughout the experiment.

### Intracellular bacterial viability studies

Epithelial cell monolayers infected at a MOI of 100 with *M. tuberculosis* strains Erdman, *Δrv3351c*, or *CMΔrv3351c*, were incubated for six hours before being washed three times with PBS and incubated for two hours in DMEM with 5% FBS and 200 μg/ml amikacin (added to inhibit the growth of remaining extracellular bacteria). The medium was then replaced with fresh DMEM. This was defined as time point 0. At 24 and 72 hpi, monolayers were washed and cells lysed with 0.1% Triton X-100. Viable bacilli were enumerated by serial dilution of each lysate in 1 x PBS/ 0.05% Tween 80 and plating on 7H11 agar. All infection assays were performed in triplicate.

### Lactate dehydrogenase (LDH) assay

For lactate dehydrogenase (LDH) release studies, A549 cells were seeded into 6-well Costar^®^ dishes at 1.0 x 10^6^ cells per well 24 hours prior to infection. On the day of infection, monolayers were washed three times with Hanks Balanced Salt Solution (HBSS) and fresh medium was added. Cells were infected in parallel with *M. tuberculosis* strains Erdman, *Δrv3351c* and *CMΔrv3351c* at a MOI of 100. All infections were performed in triplicate. For studies testing Rv3351c recombinant protein or Filipin III, 5 μg of Rv3351c or Filipin III was added to appropriate wells 30 minutes before infection with the indicated *M. tuberculosis* strain using the methods of Fine-Coulson *et al*. (9). Cells were washed and Filipin III-treated cells were maintained in 2.5 μg /mL for the duration of the experiment. Filipin III was obtained from Sigma (Sigma-Aldrich, St. Louis, MO) and reconstituted with DMSO. For Bafilomycin A1 treatment (Sigma-Aldrich, St. Louis, MO), the compound was added to a final 100 nm concentration in appropriate wells and maintained for the duration of the experiment. Host cells treated with Rv3351c, Filipin and Bafilomycin A1 were evaluated for cytotoxic effects by LDH release assay. Supernatants were sampled over a time course and filtered through PVDF membranes (0.22μm). Immediately following filtration, supernatants were assayed for LDH activity using the Cytotoxicity Detection Kit (Roche, Indianapolis, IN) (8,20). Percent LDH release was calculated using the following formula: [(Release from Strain – Background)/(Max release - Background)] x 100.

### Confocal microscopy

For confocal microscopy, A549 cells were grown as monolayers to confluency, harvested with trypsin-treatment for three minutes at 37°C, and 5 x 10^5^ cells were seeded onto sterile cover slips placed within 6-well Costar^®^ dishes. The cells were allowed to adhere for 12 hours at 37°C in 5% CO_2_ and then infected as described. Specimens were fixed at indicated time points with either 2% or 3.7% paraformaldehyde for one hour. The specimens were washed three times with 1 x PBS and the cells permeabilized for 10 minutes with 0.1% Triton X-100. The samples were then blocked for 30 minutes with PBS containing 3% BSA. For attachment studies, cells were blocked and then labeled with Alexa Fluor^®^488 or 546 phalloidin (Invitrogen, Carlsbad, CA) for 35 minutes at room temperature. For lysosomal studies, LAMP-2 monoclonal antibody (Abcam Inc., Cambridge, MA) was added to each coverslip at a dilution of 1:200 and incubated at room temperature for one hour. After washing with PBS, mouse-anti-human antibody conjugated with Texas Red was then added at a dilution of 1:500 and incubated for one hour at room temperature. Cytoskeletal staining was achieved by 35-minute incubation with Alexa Fluor^®^647 phalloidin (Invitrogen). Images were obtained with a Zeiss LSM 510 or Leica SP5 confocal microscope.

For LC3 studies, LC3b rabbit polyclonal antibody (Abcam Inc., Cambridge, MA) was added to each coverslip at a dilution of 1:200 and incubated at room temperature for one hour. After washing with PBS, goat-anti-rabbit antibody conjugated with Alexa Fluor^®^633 (Invitrogen, Carlsbad, CA) was then added at a dilution of 1:500 and incubated for 30 minutes at room temperature.

For GM1 studies, 2 μg listeriolysin O (LLO) (Abcam Inc., Cambridge, MA), Rv3351c, or Rv0097 recombinant protein were added to appropriate wells for 30 minutes at room temperature before staining. Cholera toxin subunit B conjugated to Alexa Fluor 488 (Invitrogen, Carlsbad, CA) was added at a dilution of 1:250 to all coverslips 30 minutes prior to fixation with 4% paraformaldehyde. All staining steps were carried out at 4°C. A total of 10 fields were imaged per coverslip for each experiment, incorporating approximately 50–100 host cells per field. Images were captured at 10x, 63x or 100x magnification. Two coverslips were obtained for all conditions at the designated time point for all three experimental replicates. Aggregates of cholera toxin and LC3 puncta were quantified and analyzed for size and area using ImageJ software. Numbers of aggregates were normalized to the number of host cells imaged per field to acquire an average number of aggregates per host cell/per field for each condition.

### Total cell protein extraction

At 12 and 24 hpi, cell lysates were harvested with Pierce™ IP Lysis Buffer (Pierce Biotechnology, Rockford, IL), according to the manufacturer’s instructions. For amino acid starvation conditions, cell monolayers were washed 3 times with Earle’s Balanced Salts Solution (EBSS), and 2 mL of fresh EBSS was added for 4 hours. To inhibit autophagic flux, 100 nm Bafilomycin A1 was added two hours before each time point. Supernatants were filtered through 22-μm PVDF membranes before being removed from the biosafety level 3 laboratory. Total protein concentration was determined by BCA protein quantification (ThermoFisher Scientific, CA). Protein samples were mixed with sodium dodecyl sulfate buffer (1× final concentration) and heated at 90°C for 10 minutes. Proteins were separated by electrophoresis on a 12% sodium dodecyl sulfate polyacrylamide gel. Total protein was transferred to a PVDF membrane (Bio-Rad, Hercules, CA), which was then preincubated with blocking solution (5% nonfat dry milk in Tris-buffered saline containing 0.01% Tween 20; pH, 7.4) for one hour, followed by overnight incubation with anti-LC3B or anti-ULK1 rabbit monoclonal antibodies (1:1000), or anti-GAPDH mouse monoclonal antibody (1:1000) (Cell Signaling Technology, Inc, Danvers, Massachusetts) at 4°C. After primary incubation, the membrane was washed three times with Tris-buffered saline containing 0.01% Tween 20 and incubated for one hour with secondary rabbit (Cell Signaling Technology, Inc, Danvers, Massachusetts) or goat anti-mouse HRP-conjugated antibody (Abcam Inc., Cambridge, MA). All incubations and wash steps were performed at room temperature except when otherwise stated. Cross-reactivity was visualized by using enhanced chemiluminescence (SuperSignalWestPico; Pierce Biotechnology, Rockford, IL).

### Transmission electron microscopy

For transmission electron microscopy (TEM), cells were harvested and seeded into T25 flasks at a density of 5.0 x 10^6^ cells/ml. The cell cultures were synchronized by incubation at 4°C for two hours: one hour preceding infection and one hpi. Infections were conducted as previously described. Specimens were fixed with 2.5% glutaraldehyde for one hour and then placed in phosphate buffer prior to treatment with 1% osmium tetroxide for 45 minutes and a phosphate buffer wash. An ethanol series was used to dehydrate the specimens that were then infiltrated using propylene oxide. Thorough infiltration was completed with three ratios of propylene oxide to resin (Epon-araldite). Resin recipes were based on previous protocols by Mollenhauer (50). Specimens were incubated one hour in resin followed by an exchange and overnight incubation at room temperature. After an additional resin exchange, samples were embedded and polymerized overnight at 60°C. Ultrathin sections were mounted onto copper grids and stained with 4% uranyl acetate and lead citrate. Imaging was performed using a Tecnai BioTwin electron microscope (FEI Company, Hillsboro, OR) operating at 80 or 120 kV. Digital images were captured using a 2K x 2K camera (AMT, Danvers, MA). Images were edited using Microsoft^®^ Picture Manager and Adobe^®^ Photoshop 7.0.

### Immuno-electron microscopy

For immuno-electron microscopy (IEM), cells were harvested and seeded into T25 flasks at a density of 5.0 x 10^6^ cells/ml. The cell cultures were synchronized by incubation at 4°C for two hours; one hour preceding infection and one hpi with *M. tuberculosis* strains at a MOI equivalent to 100. At the indicated time points, cells were fixed with a 1.5% paraformaldehyde/0.025% glutaraldehyde solution for one hour, and then placed in phosphate buffer. The fixed specimens were then dehydrated using a graded ethanol series. Cells were incubated in increasing ratios of ethanol toLR White embedding media as outlined by Goldsmith *et al*. (51). Samples were then allowed to incubate one hour in 100% LR White followed by a fresh exchange and overnight incubation at 4°C. The following day, specimens were incubated in fresh LR White for one hour, placed in gelatin capsules, centrifuged (1,500 x g, 5 minutes) and the blocks allowed to polymerize for 20-24 hours at 58°C. Ultrathin sections were mounted onto nickel grids and blocked with normal goat serum diluted 1:100. Each grid was incubated with a 1:500 dilution of cathepsin-L (Abcam Inc., Cambridge, MA) or LAMP-2 (Invitrogen, Carlsbad, CA) antibody for 75 minutes. Gold-conjugated secondary labels: 12-nm goat-anti-rabbit or 20-nm goat-anti-mouse IgG (Jackson ImmunoResearch) were used at a 1:20 dilution with one hour of incubation. Samples were imaged and analyzed as described for TEM.

### Intracellular quantification

Total bacteria from TEM experiments were counted within at least 20 grid fields containing 10-15 infected pneumocytes per field. The number of bacteria inside the cells was divided by the total number of bacteria within the defined grid fields to determine the percent internalization. Infections were performed in duplicate and experiments repeated three times. For confocal attachment/internalization studies, z-stack images were obtained for each field of view and external/internal bacteria quantified by slice. Infections were performed in duplicate and experiments repeated three times. For each coverslip, 10 fields of view were captured and quantified (~250 host cells counted per coverslip).

### *M. tuberculosis* membrane fraction analysis

Cultures of *M. tuberculosis* Erdman or *Δrv3351c* were grown to an OD600 of 0.8 and centrifuged at 2,000 x g for ten minutes to pellet the bacteria; pellet and supernatant fractions were stored at −80°C until further processing. Pellets were thawed and washed with equal volumes of ice-cold PBS. Washed pellets were resuspended in PBS containing complete EDTA-free protease inhibitor (Roche, Indianapolis, IN), 2 mM EDTA, and 0.5 mM PMSF, and lysed by bead beating. Lysed samples were transferred to clean microcentrifuge tubes and centrifuged at 8,000 x g for ten minutes at 4°C. Supernatants were filtered through 22 μm PVDF membranes and pellets heat-killed before being removed from biosafety level 3. Total protein concentration was determined by BCA protein quantification. Protein samples were mixed with sodium dodecyl sulfate buffer (1× final concentration) and heated at 90°C for ten minutes. Proteins were separated by electrophoresis on a 12% sodium dodecyl sulfate polyacrylamide gel. Total protein was transferred to a nitrocellulose membrane (Bio-Rad, Hercules, CA), which was then preincubated with blocking solution (5% nonfat dry milk in Tris-buffered saline containing 0.01% Tween 20; pH, 7.4) for one hour, followed by overnight incubation with anti-Rv3351c mouse monoclonal antibody (1:100), or anti-RNA polymerase beta subunit antibody (Abcam Inc., Cambridge, MA) at 4°C. After primary incubation, the membrane was washed three times with Tris-buffered saline containing 0.01% Tween 20 and incubated for one hour with secondary mouse HRP-conjugated antibody (Abcam Inc., Cambridge, MA). All incubations and wash steps were performed at room temperature unless otherwise stated. Cross-reactivity was visualized by using enhanced chemiluminescence (SuperSignalWestPico; Pierce Biotechnology, Rockford, IL)

### TNF-α, MCP-1, IL-8 ELISA

Supernatants of infected cells were assayed for cytokine levels by ELISA using the BD OptEIA™ kits for TNF-α, MCP-1, or IL-8 (BD Pharmingen, Franklin Lakes, NJ) according to manufacturers’ protocols.

### Expression and purification of recombinant proteins

Plasmids pET19b-Rv3351c and pET19b-Rv97 were generated by incorporating *rv3351c* and *rv0097c*, respectively, downstream of a hexahistidine tag. Specifically, using *M. tuberculosis* H37Rv genomic DNA as template, the *rv3351c* gene was PCR-amplified with primers 5’–GGA-ATT-CCA-TAT-GCT-GGC-GAG-CTG-CCC-GG-3’ and 5’-GAA-GAT-CTT-CAC-CGT-GGA-GTA-CGA-CGA-ACC-GGC-3’; the *rv0097c* gene was amplified with primers 5’-GGA-ATT-CCA-TAT-GAC-GCT-TAA-GGT-CAA-AGG-CGA-GGG-3’ and 5’-GAA-GAT-CTC-ATG-CCG-CGT-ATC-CCG-G-3’. Each PCR product was digested with *NdeI* and *BglII* and ligated into pET19b digested with the same enzymes. *Escherichia coli* Rosetta™ (DE3) transformants were obtained and maintained by culture in LB medium with carbenicillin (50 μg/ml) and chloramphenicol (34 μg/mL). Plasmids were confirmed free of PCR-induced mutations by DNA sequencing. For protein expression, IPTG was added to a final concentration of 0.2 mM when cultures reached OD_600_ = 0.6 and the cells were incubated 4 hours. Cells were harvested by centrifugation (6,000 x g, 15 minutes, 4°C) and pellets stored at −80°C.

Cells containing pET19b-Rv3351c were resuspended in B-PER Complete Bacterial Protein Extraction Reagent (Pierce Biotechnology, Rockford IL) with 0.5 mM PMSF, incubated for one hour with gentle rocking, and lysates clarified by centrifugation (16,000 x g, 20 minutes, 4°C). Pellets containing Rv3351c protein were washed three times with buffer (50 mM Tris–HCl (pH 8), 10 mM EDTA, 100 mM NaCl, and 0.5% Triton X-100) and solubilized in urea buffer (8 M urea, 10 mM Tris–HCl, pH 7.4) by stirring (4 hours at room temperature). Solubilize protein was obtained by collecting the supernatant after centrifugation (15,200 x g, 40 minutes, 4°C).

Cells containing pET19b-Rv97 were re-suspended in Buffer C (52) plus 0.5 mM PMSF and lysed in a One Shot Disrupter (Constant Systems, Inc.). After centrifugation (21,000 x g, 15 minutes), the pellet was suspended in Buffer C plus 6M urea.

Nickel-NTA chromatography was used to purify the recombinant 6His-tagged Rv3351c and Rv0097 proteins. The urea-solubilized proteins were applied to Ni2+-charged HiTrap columns pre-equilibrated with urea phosphate buffer (8 M urea in 20 mM sodium phosphate, pH 7.4). The column was washed with the same buffer prior to eluting proteins with a linear gradient of imidazole (10–500 mM) in urea phosphate buffer. Proteins were renatured by stepwise dialysis at 4°C against 5 mM Tris (pH 7.4) containing decreasing concentrations of urea: 6, 4, 2, 1, 0.5, 0.01, and 0 M, respectively.

### Human clinical serum ELISA

Two or 4 ng of Rv3351c protein was immobilized on a microtiter plate (Nunc,ThermoFisher Scientific, CA) overnight at 4°C. The next day, the plate was washed three times with phosphate-buffered saline (pH 7.4) containing 0.05% Tween 20 (PBS-Tween20) and blocked for 30 minutes with 5% bovine serum albumin diluted in PBS-Tween20. The plate was washed three times with PBS-Tween20 before incubation for one hour at room temperature with human sera diluted 1:20, 1:50, 1:100, and 1:200 in PBS from patients determined to have culture-confirmed active or latent infection or from uninfected controls (33). After washing, a 30-minute incubation with goat anti-human HRP-conjugated secondary antibody (Invitrogen, Carlsbad, CA) was performed prior to incubation with 1-Step™ Ultra TMB-ELISA Substrate Solution (ThermoFisher Scientific, CA) for color development, which was measured spectrophotometrically at 405 nm.

### Mouse aerosol infections

BALB/c mice were obtained from Charles River Laboratory. BALB/c mice were challenged with 1X10^5^ CFU *M. tuberculosis* Erdman or the *CMΔrv3351c* complemented strain in 0.025 ml PBS via the intratracheal route (53). Four mice per group were sacrificed 24 hpi to confirm actual exposure dose in the lungs. All mice were monitored daily over the course of the three-week study for malaise or other adverse reactions. After euthanasia, portions of tissues including lungs, livers, and spleens were harvested, fixed in 10% neutral-buffered formalin, and processed for histopathology. The histologic slides were scored by a pathologist blinded to the identity of the individual experimental samples (54). In addition, portions of the collected tissues were placed in PBS with 0.05% Tween 80 and homogenized. The homogenates were serially-diluted and plated onto 7H10gtADC agar plates and incubated at 37°C for three weeks prior to colony count assessments. All mice were monitored daily over the course of the 21-day study for weight loss, malaise or other adverse reactions. Mice were euthanized based on a quantitative assessment of disease endpoints.

### Thin layer chromatography of lipids extracted from A549 cells

Total glycosphingolipid extracts were prepared from A549 cells as previously described {{*PMID: 24026681}}. Thin layer chromatography overlay technique was conducted as previously described {{*PMID: 1325706, PMID: 1601891, PMID: 29926422}}. Briefly, 2.5 μL of A549 lipid extracts or neutral glycosphingolipid, ganglioside, and lactosylceramide-derived glycosphingolipid standards were spotted onto a silica gel 60 glass plate (with fluorescent indicator) and the plate was developed using a mobile phase solvent composed of chloroform:methanol:0.2% aqueous CaCh::60/40/10 (v/v/v). After drying, the plates were sequentially dipped in hexane followed by 0.005% polyisobutylmethacrylate (Aldrich) in hexane for 30 s each. The plate was allowed to air dry and was then wetted by spray with PBS. The sprayed plate reveals the position of lipid migration as white spots against a grey background due to differential wetting of the hydrophobic components (see Supplement Figure 2). After wetting with PBS, the TLC plate was blocked to minimize non-specific binding by overlaying a solution of PBS containing 1.0% BSA and incubating at room temperature for 1h. During the blocking step, 50 μg of recombinant Rv3351c protein and anti-6xHIS antibody conjugated to horseradish peroxidase (HRP, 1:500; Abcam Inc., Cambridge, MA) were mixed together in PBS containing 0.5% BSA. The resulting pre-formed protein-antibody complexes were then overlaid onto the blocked TLC plate and incubated for 1 hour at room temperature. The plate was then washed three times for 15 minutes in PBS and subsequently incubated for 10 minutes at room temperature with 0.5mg/ml diaminobenzidine and 0.003% hydrogen peroxide for detection of HRP enzyme activity. In parallel, an identical plate was prepared for detection of glycolipids by orcinol staining.

### Statistical analysis

Statistical significance was determined by ANOVA and Tukey’s HSD post-hoc comparison using GraphPad Prism 8.0 (GraphPad Software, San Diego, CA, USA). Animal organ mycobacterial burden comparisons were made using Student’s t test and the Mann-Whitney U test within the program GraphPad Prism 5.0 (GraphPad Software, San Diego, CA, USA). Animal survival curves were analyzed using the Kaplan Meier estimator in GraphPad Prism 5.0.

## Supporting information

**Supplemental Figure 1.** Rv3351c binds hydrophobic glycolipids and hydrophobic components of the lipids extracted from A549 cells. Lipid extracts were spotted and resolved on thin-layer silica plates along with three sets of well-characterized sets of glycosphingolipid standards. One plate was sprayed with Orcinol (*left panel*) to detect glycosylated lipids. An identical plate was prepared, developed, and coated with plastic for probing with recombinant Rv3351c (*middle panel*). The plastic-coated plate was sprayed with PBS, revealing the presence of hydrophobic compounds based on differential wetting, including cholesterols, fatty acids, and glycolipids (*right panel*) before being probed with Rv3351c. Standards, with cartoon representations, included in each of the indicated sets and their migration positions are shown on the left. The N Standard set included glucosyl-ceramide (GlcCer), lactosyl-ceramide (LacCer), globotriaosyl-ceramide (Gb3Cer), and globotetraosylceramide (Gb4Cer), from fastest to slowest migrating. The G Standard set included asialo-gangliotetraosylceramide (Gg4Cer) as well as GM1, GD1a, GD1b, and GT1b gangliosides. The L Standard set included lactosyl-ceramide LacCer, which is also in the N-Standard set, as well as GM3, and GD3 ganglioside. Orcinol staining indicates that the major glycolipids in the A549 extract have chromatographic mobility consistent with the assignment as asialo-gangliotriosylceramide Gg3Cer (*a*), GM2 (*b*), and GD1a (*c*), as has been recently described by high-sensitivity LC-MS/MS analysis {{*PMID: 31717732}}. Rv3351c binds to a major orcinol-negative, hydrophobic component in the lipid extract of A549 cells, most likely cholesterol, and another minor orcinol-negative component (both indicated by asterisk). Rv3351c also binds to orcinol-positive glycolipids extracted from A549 cells (indicated by arrowheads). The major glycolipids bound by Rv3351c do not migrate with any of the standard set lipids and the standard set lipids bind Rv3351c poorly, despite being presented at equal or higher density to A549 components on the silica plates. The glycolipid components of the A549 extract exhibit greater potency for binding Rv3351c than the standard lipids on the silica plate, despite being present at lower amounts (compare orcinol staining). Even for component c, assigned as GD1a which is present in the G Standard set, Rv3351c preferred the A549 lipid over the standard, suggesting that both the glycan and the unique ceramide composition of the A549 lipid contribute to binding.

**Supplemental Figure 2.** (A) Monoclonal antibodies made against *E. coli* recombinant Rv3351c recognize a protein of the expected size for Rv3351c in in the membrane fraction and to a lesser extent in the cytoplasmic fraction of *M. tuberculosis* strain Erdman cell lysates by western blot. (B) Anti-RNA polymerase β subunit antibody served as a control for the cytoplasmic fraction which detects a band of approximately 140 kDa. S = supernatant; C = cytoplasmic; M = membrane fraction.

**Supplemental Figure 3.** Lung sections from BALB/c mice intranasally-infected with *Δrv3351c* or bacilli from the complemented strain, *CMΔrv3351c* show similar infiltration in BALB/c mouse lungs 21 days post-infection. Healthy lung tissue is visible on day 1. Arrows highlight areas of lymphocyte infiltration on day 21 post-infection. The intratracheal dose delivered was 1X10^5^ CFU in 0.025 ml PBS. Organs were harvested post-euthanasia and tissue sections stained with hematoxylin and eosin as described.

## Acknowledgements

This work was supported in part by research grants from the University of Georgia Faculty of Infectious Diseases/Southeastern Center for Emerging Biologic Threats (F.Q.), and the American Lung Association (R.K.). M.M. current affiliation is Deloitte Consulting LLP, Atlanta, Georgia.

## Author contributions

### Conceptualization

Megan Prescott, Kari Fine-Coulson, Rebecca Pavlicek, Frederick Quinn

### Data curation

Megan Prescott, Kari Fine-Coulson, Frederick Quinn

### Formal analysis

Megan Prescott, Kari Fine-Coulson, Frederick Quinn

### Funding acquisition

Fred Quinn

### Investigation

Megan Prescott, Kari Fine-Coulson, Maureen Metcalfe, Tuhina Gupta, Michelle Dookwah, Rebecca Pavlicek, Barbara Reaves, Russell Karls, Frederick Quinn

### Methodology

Megan Prescott, Kari Fine-Coulson, Maureen Metcalfe, Tuhina Gupta, Michelle Dookwah, Rebecca Pavlicek, Barbara Reaves, Russell Karls, Frederick Quinn

### Project administration

Frederick Quinn

### Resources

Megan Prescott, Kari Fine-Coulson, Maureen Metcalfe, Tuhina Gupta, Michelle Dookwah, Rebecca Pavlicek, Barbara Reaves, Russell Karls, Frederick Quinn

### Software

Megan Prescott, Kari Fine-Coulson, Barbara Reaves, Frederick Quinn

### Supervision

Frederick Quinn

### Validation

Megan Prescott, Kari Fine-Coulson, Frederick Quinn

### Writing – original draft

Megan Prescott, Kari Fine-Coulson, Frederick Quinn

### Writing – review and editing

Megan Prescott, Kari Fine-Coulson, Maureen Metcalfe, Tuhina Gupta, Michelle Dookwah, Rebecca Pavlicek, Barbara Reaves, Russell Karls, Frederick Quinn

## References

1. World Health Organization: Global Tuberculosis Report (2018). https://www.publichealthupdate.com/global-tuberculosis-report-2018-world-health-organization

2. Hauck, F., Neese, B.H., Panchal, A.S. & El-Amin, W. Identification and management of latent tuberculosis infection. Am. Fam. Physician 79, 879–886 (2009).

3. Armstrong, J.A. & Hart, D. Response of cultured macrophages to *Mycobacterium tuberculosis*, with observations on fusion of lysosomes with phagosomes. J. Exp. Med. 134, 713–740 (1971).

4. Joshi, N., Walter, J.M., & Misharin, A.V. Alveolar Macrophages. Cell Immunol. 330:86–90 (2018).

5. Ryndak, M.B., Singh. K.K., Peng, Z., & Laal, S. Transcriptional profile of *Mycobacterium tuberculosis* replicating in type II alveolar epithelial cells. PLoS One. 10(4), e0123745 (2015).

6. Ryndak, M.B. & Laal, S. *Mycobacterium tuberculosis* primary infection and dissemination: A critical role for alveolar epithelial cells. Front. Cell. Infect. Microbiol. 9, 299 (2019).

7. Scordo, J.M., Knoell, D.L. & Torrelles, J.B. Alveolar epithelial cells in *Mycobacterium tuberculosis* infection: Active players or innocent bystanders? J. Innate Immun. 8, 3–14 (2016).

8. Pavlicek, R.L., Fine-Coulson, K., Gupta, T., Quinn, F.D., Posey, J.E., Willby, M., Castro-Garza, J. & Karls, R.K. Rv3351c, a *Mycobacterium tuberculosis* gene that affects bacterial growth and alveolar epithelial cell viability. Can. J. Microbiol. 61, 938–947 (2015).

9. Fine-Coulson, K., Reaves, B.J., Karls, R.K. & Quinn, F.D. The role of lipid raft aggregation in the infection of type II pneumocytes by *Mycobacterium tuberculosis*. PLoS One 7(9):e45028 (2012).

10. Birkness, K.A., Deslauriers, M., Bartlett, J.H., White, E.H., King, C.H. & Quinn, F.D. An *in vitro* tissue culture bilayer model to examine early events in *Mycobacterium tuberculosis* infection. Infect. Immun. 67, 653–658 (1999).

11. Bermudez, L.E. & Goodman, J. *Mycobacterium tuberculosis* invades and replicates within type II alveolar cells. Infect. Immun. 64, 1400–1406 (1996).

12. Mehta, P.K., King, C.H., White, E.H., Murtagh, J.J. & Quinn, F.D. Comparison of *in vitro* models for the study of *Mycobacterium tuberculosis* invasion and intracellular replication. Infect. Immun. 64, 2673–2679 (1996).

13. Pethe, K., Alonso, S., Biet, F., Delogu, G., Brennan, M.J., Locht, C. & Menozzi, F.D. The heparin-binding haemagglutinin of *M. tuberculosis* is required for extrapulmonary dissemination. Nature 412, 190–194 (2001).

14. Hsu, T., Hingley-Wilson, S.M., Chen, B., Chen, M., Dai, A.Z., Morin, P.M., Marks, C.B., Padiyar, J., Goulding, C., Gingery, M., Eisenberg, D., Russell, R.G., Derrick, S.C., Collins, F.M., Morris, S.L., King, C.H. & Jacobs, W.R. Jr. The primary mechanism of attenuation of bacillus Calmette-Guérin is a loss of secreted lytic function required for invasion of lung interstitial tissue. Proc. Natl. Acad. Sci. U. S. A. 100, 12420–12425 (2003).

15. Chitale, S., Ehrt, S., Kawamura, I., Fujimura, T., Shimono, N., Anand, N., Lu, S., Cohen-Gould, L. & Riley, L.W. Recombinant *Mycobacterium tuberculosis* protein associated with mammalian cell entry. Cell. Microbiol. 3, 247–254 (2001).

16. Fine, K.L., Metcalfe, M.G., White, E., Virji, M., Karls, R.K. & Quinn, F.D. Involvement of the autophagy pathway in trafficking of *Mycobacterium tuberculosis* bacilli through cultured human type II epithelial cells. Cell. Microbiol. 14, 1402–1414 (2012).

17. Racanelli, A.C., Kikkers, S.A., Choi, A.M. K. & Cloonan, S.M. Autophagy and inflammation in chronic respiratory disease. Autophagy vol. 14 221–232 (2018).

18. Guo, X.G., Ji, T.X., Xia, Y. & Ma, Y.Y. Autophagy protects type II alveolar epithelial cells from *Mycobacterium tuberculosis* infection. Biochem. Biophys. Res. Commun. 432, 308–313 (2013).

19. McDonough, K.A. & Kress, Y. Cytotoxicity for lung epithelial cells is a virulence-associated phenotype of *Mycobacterium tuberculosis*. Infect. Immun. 63, 4802–4811 (1995).

20. Dobos, K.M., Spotts, E.A., Quinn, F.D. & King, C.H. Necrosis of lung epithelial cells during infection with *Mycobacterium tuberculosis* is preceded by cell permeation. Infect. Immun. 68, 6300–6310 (2000).

21. Danelishvili, L., McGarvey, J., Li, Y.J. & Bermudez, L.E. *Mycobacterium tuberculosis* infection causes different levels of apoptosis and necrosis in human macrophages and alveolar epithelial cells. Cell. Microbiol. 5, 649–660 (2003).

22. Keane, J., Remold, H.G. & Kornfeld, H. Virulent *Mycobacterium tuberculosis* strains evade apoptosis of infected alveolar macrophages. J. Immunol. 164, 2016–2020 (2000).

23. García-Pérez, B.E., Mondragón-Flores, R. & Luna-Herrera, J. Internalization of *Mycobacterium tuberculosis* by macropinocytosis in non-phagocytic cells. Microb. Pathog. 35, 49–55 (2003).

24. Muñoz, S., Rivas-Santiago, B. & Enciso, J.A. *Mycobacterium tuberculosis* entry into mast cells through cholesterol-rich membrane microdomains. Scand. J. Immunol. 70, 256–263 (2009).

25. Carlsson, S.R. & Simonsen, A. Membrane dynamics in autophagosome biogenesis. J. Cell Sci. 128, 193–205 (2015).

26. Schleimer, R.P., Kato, A., Kern, R., Kuperman, D. & Avila, P.C. Epithelium: At the interface of innate and adaptive immune responses. J. Allergy Clin. Immunol. 120, 1279–1284 (2007).

27. Olsen, A., Chen, Y., Ji, Q., Zhu, G., De Silva, A.D., Vilchèze, C., Weisbrod, T., Li, W., Xu, J., Larsen, M., Zhang, J., Porcelli, S.A., Jacobs, W.R. Jr. & Chan, J. Targeting *Mycobacterium tuberculosis* tumor necrosis factor alpha-downregulating genes for the development of antituberculous vaccines. MBio 7, (2016).

28. Chuquimia, O.D., Petursdottir, D.H., Rahman, M.J., Hart, K., Singh, M. & Fernández, C., The role of alveolar epithelial cells in initiating and shaping pulmonary immune responses: Communication between innate and adaptive immune systems. PLoS One 7, (2012).

29. Wickremasinghe, M.I., Thomas, L.H. & Friedland, J.S. Pulmonary epithelial cells are a source of IL-8 in the response to *Mycobacterium tuberculosis:* essential role of IL-1 from infected monocytes in a NF-kappa B-dependent network. J. Immunol. 163, 3936–47 (1999).

30. Li, Y., Wang, Y. & Liu, X. The role of airway epithelial cells in response to Mycobacteria infection. Clin. Dev. Immunol. 2012, 11 (2012).

31. Shin, A.R., Lee, K.S., Lee, J.S., Kim, S.Y., Song, C.H., Jung, S.B., Yang, C.S., Jo, E.K., Park, J.K., Paik, T.H. & Kim, H.J. *Mycobacterium tuberculosis* HBHA protein reacts strongly with the serum immunoglobulin M of tuberculosis patients. Clin. Vaccine Immunol. 13, 869–875 (2006).

32. Stylianou, E., Diogo, G.R., Pepponi, I., van Dolleweerd, C., Arias, M.A., Locht, C., Rider, C.C., Sibley, L., Cutting, S.M., Loxley, A., Ma, J.K. & Reljic, R. Mucosal delivery of antigen-coated nanoparticles to lungs confers protective immunity against tuberculosis infection in mice. Eur. J. Immunol. 44, 440–449 (2014).

33. Castro-Garza, J., García-Jacobo, P., Rivera-Morales, L.G., Quinn, F.D., Barber, J., Karls, R., Haas, D., Helms, S., Gupta, T., Blumberg, H., Tapia, J., Luna-Cruz, I., Rendon, A., Vargas-Villarrea, J., Vera-Cabrera, L. & Rodríguez-Padilla, C. Detection of anti-HspX antibodies and HspX protein in patient sera for the identification of recent latent infection by Mycobacterium tuberculosis. PLoS One 12, (2017).

34. Maier, B.B., Hladik, A., Lakovits, K., Korosec, A., Martins, R., Kral, J.B., Mesteri, I., Strobl, B., Müller, M., Kalinke, U., Merad, M. & Knapp, S. Type I interferon promotes alveolar epithelial type II cell survival during pulmonary *Streptococcus pneumoniae* infection and sterile lung injury in mice. Eur J Immunol. 46(9), 2175–86 (2016).

35. Thomas, R. & Brooks, T. Attachment of *Yersinia pestis* to human respiratory cell lines is inhibited by certain oligosaccharides. J. Med. Microbiol. 55, 309–315 (2006).

36. Thomas, R. J. & Brooks, T. J. Oligosaccharide receptor mimics inhibit *Legionella pneumophila* attachment to human respiratory epithelial cells. Microb. Pathog. 36, 83–92 (2004).

37. Debbabi, H., Ghosh, S., Kamath, A.B., Alt, J., Demello, D.E., Dunsmore, S. & Behar, S.M. Primary type II alveolar epithelial cells present microbial antigens to antigen-specific CD4 ^+^ T cells. Am. J. Physiol. Cell. Mol. Physiol. 289, L274–L279 (2005).

38. Manabe, Y.C., Dannenberg, A.M., Jr., Tyagi, S.K., Hatem, C.L., Yoder, M., Woolwine, S.C., Zook, B.C., Pitt, M.L. & Bishai, W.R. Different strains of *Mycobacterium tuberculosis* cause various spectrums of disease in the rabbit model of tuberculosis. Infect. Immun. 71, 6004–6011 (2003).

39. McDonough, K. A., Kress, Y. & Bloom, B. R. Pathogenesis of tuberculosis: Interaction of *Mycobacterium tuberculosis* with macrophages. Infect. Immun. 61, 2763–2773 (1993).

40. Sanjuan, M.A., Dillon, C.P., Tait, S.W., Moshiach, S., Dorsey, F., Connell, S., Komatsu, M., Tanaka, K., Cleveland, J.L., Withoff, S. & Green, D.R. Toll-like receptor signalling in macrophages links the autophagy pathway to phagocytosis. Nature 450, 1253–1257 (2007).

41. Lerena, M. C. & Colombo, M. I. *Mycobacterium marinum* induces a marked LC3 recruitment to its containing phagosome that depends on a functional ESX-1 secretion system. Cell. Microbiol. 13, 814–835 (2011).

42. Köster, S., Upadhyay, S., Chandra, P., Papavinasasundaram, K., Yang, G., Hassan, A., Grigsby, S.J., Mittal, E., Park, H.S., Jones, V., Hsu, F.F., Jackson, M., Sassetti, C.M. & Philips, J.A. *Mycobacterium tuberculosis* is protected from NADPH oxidase and LC3-associated phagocytosis by the LCP protein CpsA. Proc. Natl. Acad. Sci. U. S. A. 114, E8711–E8720 (2017).

43. Schille, S., Crauwels, P., Bohn, R., Bagola, K., Walther, P. & van Zandbergen, G. LC3-associated phagocytosis in microbial pathogenesis. Intl. J. Med. Microbiol. 308, 228–236 (2018).

44. Heckmann, B. L. & Green, D. R. LC3-associated phagocytosis at a glance. J. Cell Sci. 132 (2019).

45. Martín-Martín, A.I., Vizcaíno, N. & Fernández-Lago, L. Cholesterol, ganglioside GM1 and class A scavenger receptor contribute to infection by *Brucella ovis* and *Brucella canis* in murine macrophages. Microbes Infect. 12, 246–251 (2010).

46. Hatayama, S., Shimohata, T., Amano, S., Kido, J., Nguyen, A.Q., Sato, Y., Kanda, Y., Tentaku, A., Fukushima, S., Nakahashi, M., Uebanso, T., Mawatari, K., & Takahashi, A. Cellular tight junctions prevent effective *Campylobacter jejuni* invasion and inflammatory barrier disruption promoting bacterial invasion from lateral membrane in polarized intestinal epithelial cells. Front. Cell. Infect. Microbiol. 8, 15 (2018).

47. Braunstein, M., Bardarov, S. S. & Jacobs, W. R. Genetic methods for deciphering virulence determinants of *Mycobacterium tuberculosis*. Meth. Enzymol. 358, 67–99 (2002).

48. Kong, D. & Kunimoto, D. Y. Secretion of human interleukin 2 by recombinant *Mycobacterium bovis* BCG. Infect. Immun. 63, 799–803 (1995).

49. Wagner, D., Sangari, F. J., Kim, S., Petrofsky, M. & Bermudez, L. E. *Mycobacterium avium* infection of macrophages results in progressive suppression of interleukin-12 production *in vitro* and *in vivo*. J. Leukoc. Biol. 71, 80–8 (2002).

50. Mollenhauer, H. H. Plastic embeffing mixtures for use in electron microscopy. Stain Technol. 39, 111–4 (1964).

51. Goldsmith, C. S., Elliott, L. H., Peters, C. J. & Zaki, S. R. Ultrastructural characteristics of Sin Nombre virus, causative agent of hantavirus pulmonary syndrome. Arch. Virol. 140, 2107–2122 (1995).

52. Rodrigue, S., Brodeur, J., Jacques, P.E., Gervais, A.L., Brzezinski, R. & Gaudreau, L. Identification of mycobacterial sigma factor binding sites by chromatin immunoprecipitation assays. J. Bacteriol. 189, 1505–1513 (2007).

53. Perdomo, C., Zedler, U., Kühl, A.A., Lozza, L., Saikali, P., Sander, L.E., Vogelzang, A., Kaufmann, S.H. & Kupz, A. Mucosal BCG vaccination induces protective lung-resident memory T cell populations against tuberculosis. MBio 7, (2016).

54. Chen, Z., Gupta, T., Xu, P., Phan, S., Pickar, A., Yau, W., Karls, R.K., Quinn, F.D., Sakamoto, K. & He, B. Efficacy of parainfluenza virus 5 (PIV5)-based tuberculosis vaccines in mice. Vaccine 33, 7217–7224 (2015).

